# Post-transcriptional RNA stabilization of telomere-proximal RNAs FRG2, DBET, D4Z4 at human 4q35 in response to genotoxic stress and D4Z4 macrosatellite repeat length

**DOI:** 10.1101/2024.03.18.585486

**Authors:** Valentina Salsi, Francesca Losi, Monica Salani, Paul D. Kaufman, Rossella Tupler

**Affiliations:** Department of Biomedical, Metabolic and Neural Sciences, University of Modena and Reggio Emilia, 41125, Modena, Italy; Center for Human Genetic Research, Massachusetts General Hospital Research Institute and Department of Neurology, Harvard Medical School, 185 Cambridge Street, Boston, Massachusetts 02114 USA; Department of Molecular, Cell and Cancer Biology, University of Massachusetts Medical School, Worcester, MA 01605 USA

**Keywords:** D4Z4 chromatin signature, DNA damage, epigenetic regulation, FSHD

## Abstract

**Background:** Reduced copy number of the D4Z4 macrosatellite at human chromosome 4q35 is associated with facioscapulohumeral muscular dystrophy (FSHD). A pervasive idea is that chromatin alterations at the 4q35 locus following D4Z4 repeat unit deletion lead to disease via inappropriate expression of nearby genes. Here, we sought to analyze transcription and chromatin characteristics across 4q35 and how these are affected by D4Z4 deletions and exogenous stresses.

**Results:** We found that the 4q subtelomere is subdivided into discrete domains, each with characteristic chromatin features associated with distinct gene expression profiles. Centromere-proximal genes within 4q35 (*ANT1*, *FAT1* and *FRG1)* display active histone marks at their promoters. In contrast, poised or repressed markings are present at telomere-proximal loci including *FRG2, DBE-T* and *D4Z4*. We discovered that these discrete domains undergo region-specific chromatin changes upon treatment with chromatin enzyme inhibitors or genotoxic drugs. We demonstrated that the 4q35 telomere-proximal *FRG2, DBE-T* and *D4Z4*-derived transcripts are induced upon DNA damage to levels inversely correlated with the D4Z4 repeat number, are stabilized through post-transcriptional mechanisms upon DNA damage, and are bound to chromatin.

**Conclusion:** Our study reveals unforeseen biochemical features of RNAs from clustered transcription units within the 4q35 subtelomere. Specifically, the *FRG2, DBE-T* and *D4Z4*-derived transcripts are chromatin-associated and are stabilized post-transcriptionally after induction by genotoxic stress. Remarkably, the extent of this response is modulated by the copy number of the D4Z4 repeats, raising new hypotheses about their regulation and function in human biology and disease.

## INTRODUCTION

Repetitive DNA sequences occur in multiple copies and comprise over 50% of the human genome [1,2]. DNA repeats can be categorized based on their size and copy number: high-frequency repeats, also known as satellite DNA sequences, are found in various loci, including pericentromeric, subtelomeric, and interstitial regions. Satellites typically form constitutive blocks of heterochromatin, notably at telomeres, centromeres and at the short arms of acrocentric chromosomes. Although most satellites (∼89.5%) are located within repressive chromatin domains, multiple studies have found that these elements have significant impacts on evolution, genetic variation and gene expression regulation [3,4].

Notably, loci proximal to telomeres are particularly prone to regulation by repetitive elements. For example, reporter genes inserted next to telomeres are silenced, a phenomenon known as telomere position effect (TPE), and silencing is further increased by upon elongation of telomeric repeats[5–7]. Several observations indicate that telomeric and subtelomeric repeats can synergically regulate transcription of nearby genes through long-distance interactions, in a manner proportional to repeat lengths. Alterations in these interactions have been implicated in a wide spectrum of human diseases [5,8], including Facioscapulohumeral Muscular Dystrophy (FSHD) (MIM 158900). FSHD is linked to deletions that reduce the copy number of the tandemly-arrayed D4Z4 macrosatellites at the 4q35 subtelomere (25-50 kb from the telomere) [9,10] (Figure 1A).

**Figure 1.**
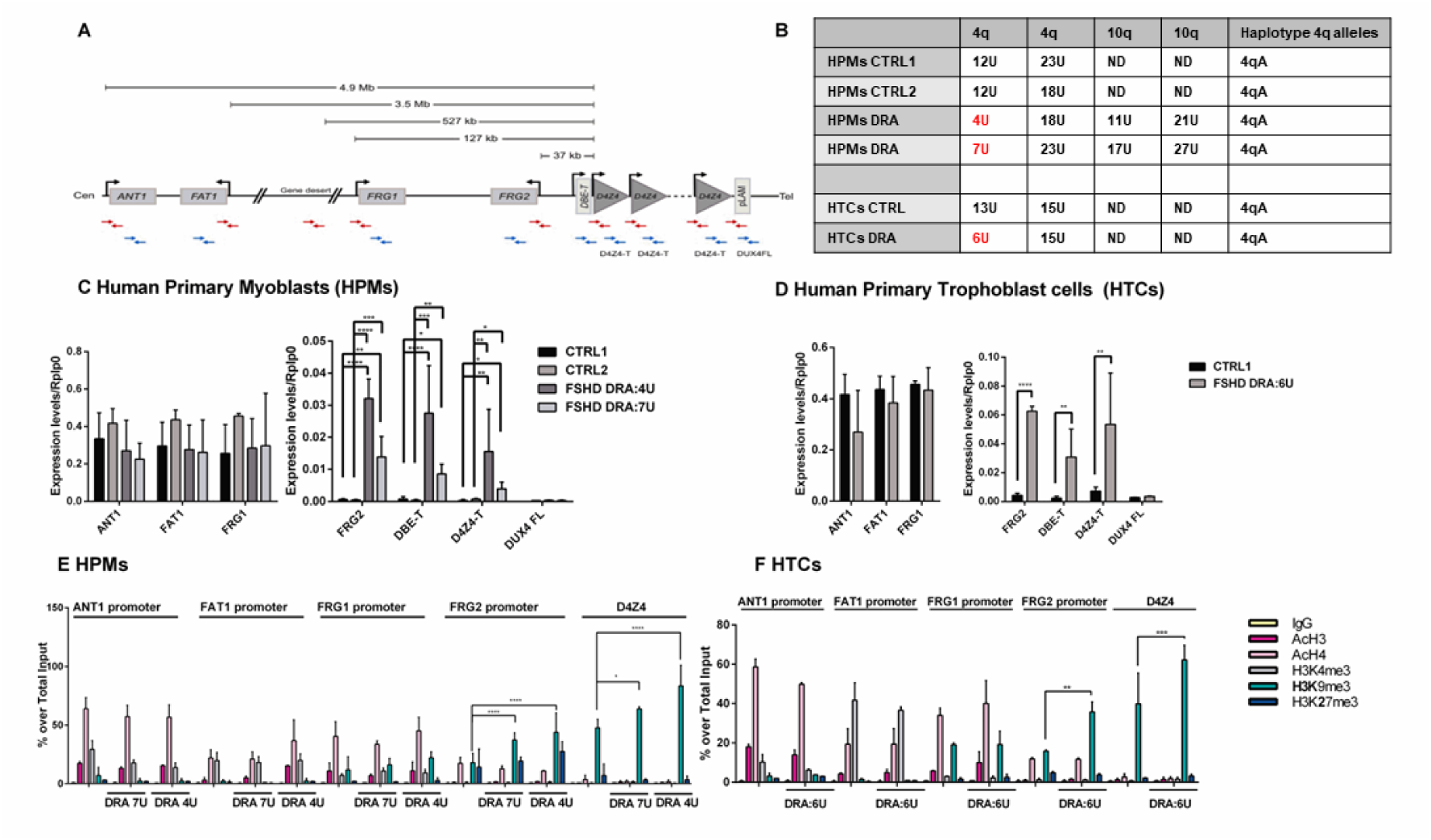
4q35 genes expression and epigenetic profile: A) Schematic representation of the chromosome 4q35 showing physical distances between *ANT1*, *FAT1*, *FRG1* and *FRG2* genes and the D4Z4 macrosatellite within the AF146191-U85056 contig, based on GenBank entry U85056.1. The positions of oligonucleotides used in ChIP experiments (red) and qPCR (blue) are shown. B) Table showing the sizes of the two 4q35 and 10q26 alleles in the selected human primary cells used in this paper, together with the 4q-ter (4qA) haplotype. Control human trophoblast cells (HTCs) and human primary myoblasts (HPMs) cells carry normal-sized 4q alleles (≥10 D4Z4 repeat units), whereas FSHD-derived HTCs and HPMs bear a reduced D4Z4 allele (DRA), i.e. < 8 D4Z4 repeat units (U=Units); C-D) RT q-PCR quantification of *ANT1*, *FAT1*, *FRG1*, *FRG2*, *DBE-T*, *D4Z4-T* and *DUX4FL* mRNAs in (C) human primary myoblasts (HPMs) and (D) human trophoblast cells (HTCs). Data were normalized using *RPLP0* as a reference mRNA. E-F) Chromatin immunoprecipitation (ChIP) analysis performed in (E) HPMs and (F) HTCs. IPs were performed using the indicated antibodies recognizing H3K4me3, H3K9me3, H3K27me3 and pan-acetylated Histone 3 and 4 (AcH3 and AcH4), or a non-specific control (IgG), followed by qPCR amplification using primers described in Fig.1A. Data are displayed as the percent enrichment for each antibody over total input chromatin. Experiments were done in triplicate and analyzed using two-way Anova statistical tests. Asterisks indicate the statistical significance of data obtained in DRA cells compared to control cells for each antibody, as follows: * 0.05<p-value<0.01; ** 0.01<p-value<0.001; *** 0.001<p-value<0.0001; **** p-value<0.0001.

The D4Z4 repeat is extremely polymorphic in the general population [11]. Earlier studies hinted at a broad distribution of these elements, since a tandemly arrayed D4Z4 macrosatellite with 98% identity to the 4q35 array is also present at subtelomere 10q26 [12,13]. By performing comprehensive bioinformatic analyses using the T2T assembly [14,15] and a collection of 86 genome assemblies from the Human Pan-Genome project [15], we have recently defined the exact number and arrangements of D4Z4-like repeat elements in the human genome [16], detecting huge inter- and intra-individual variation. Our analyses uncovered hundreds of D4Z4-like elements, which together comprise from 0.7 to 1.5 Mb of DNA, depending on the individual. We confirmed that in addition to the tandemly-arrayed D4Z4 macrosatellites at 4q35 and 10q26, incomplete D4Z4-like (D4Z4-l) sequences annotated as Beta satellites/Sau3A DNAs (Bsat) are localized at the centromeric satellite arrays surrounding rDNA repeats on the short arms of all acrocentric chromosomes and at the centromere of chromosome 1 [17–19]. At these loci, D4Z4-l sequences are not arrayed, and most of them are incomplete, lacking the 5’ portion of the D4Z4 repeat (1-800 nt) which contains regulatory regions [16]. The role and the transcriptional activity of these sequences remains obscure. However, we note that both rDNA repeats and centromeric satellites are spatially organized within nucleolus-associated domains (NADs) which are globally associated with repressive chromatin states, low gene density and low transcriptional activity [20–22] raising the idea that dispersed macrosatellites might also be involved in three-dimensional regulation of nuclear functions.

Intriguingly, the 10q26 and the 4q35 loci are notably different in their subnuclear positioning. The 4q but not the 10q subtelomere is frequently associated with the nuclear periphery. 4q35 is thus categorized as a Lamin-Associated Domain (LAD), and is thus part of a major class of generally silenced heterochromatin [20,23–25]. This observation led to the hypothesis that FSHD pathogenesis might be affected by the three-dimensional localization of the 4q35 locus [26–28]. This idea was attractive because it could explain regulation by the number of *D4Z4* elements present.

Studies investigating D4Z4 chromatin have detected multiple repressive modifications, including DNA methylation [29–35], di/trimethylation of histone H3 at lysine 9 (H3K9me2/3) [36–38], trimethylation of histone H3 at lysine 27 (H3K27me3) [39], and association of the related chromatin proteins CBX3/HP1γ and EZH2 [39,40]. Moreover, D4Z4 recruits a multi-protein repressor complex, the <u>D</u>4Z4 <u>R</u>ecognition <u>C</u>omplex (DRC), whose removal is associated with increased expression of 4q35 genes [41]. In the current patho-physiological model for FSHD [42], reduction of the D4Z4 array causes a loss of repressive histone modifications and the ectopic expression of *DUX4*, a retrogene present in each D4Z4 unit [43]. Functional, full-length DUX4 transcripts are produced from the last D4Z4 unit when they are stabilized in individuals carrying a permissive 4qA haplotype that includes a poly-adenylation signal (pLAM) near the most telomere-proximal D4Z4 unit [44]. For this reason, the majority of studies of epigenetic mechanisms in FSHD focus on the D4Z4 last repeat [45].

Here, we characterize dynamic regulatory events across the 4q35 locus. Different patterns of histone modifications characterize different chromatin domains, which correlate with the differential transcriptional responses to genotoxic stress. Additionally, drugs targeting histone modifying enzymes induce transcriptional derepression spanning the entire locus. In contrast, DNA damaging agents induce post-transcriptional stabilization of only the most telomere-proximal 4q35 transcripts, which are chromatin-associated. These latter properties are affected by the size of the subtelomeric D4Z4 array. Collectively, our results highlight that 4q35 constitutes a multipartite genomic locus providing distinct modes of regulation in response to external cues. The responses correlate with D4Z4 size thus providing additional elements to define the biological role of subtelomeric repeats and how they can be associated to anomalous responses leading to disease.

## RESULTS

### Chromatin and transcription analysis of the 4q35 region reveals distinct functional domains

To investigate the mechanism(s) by which the D4Z4 macrosatellite array affects the transcriptional regulation of the 4q35 locus (Figure 1A), we measured RNA levels of genes located at different distances from the D4Z4 array. We analyzed human primary myoblasts (HPMs) and trophoblast derived cells (HTCs) obtained from FSHD subjects heterozygous for a D4Z4 reduced allele (DRA) and matched controls (Figure 1B) [46]. On the centromere-proximal side of 4q35, we analyzed RNAs from these protein-coding genes (Figure 1A): *ANT1 (Adenine Nucleotide Translcator1;* known also as *Solute carrier family 25 member 4 SLC25A4)*, *FAT1 (FAT atypical cadherin 1*), and *FRG1 (FSHD region gene 1)*, which are are positioned at 4.9 Mb, 3.5 Mb and 127 kb from the D4Z4 array, respectively. We also analyzed telomere-proximal RNAs, including *FRG2 (FSHD region gene 2;* 37 kb from the array), and *DBE-T (D4Z4 Binding Element-Transcript)*, a lncRNA transcribed from the 5’ end of the D4Z4 array. We also analyzed two additional transcripts derived from D4Z4: DUX4 exon1 (Ex1)-containing transcripts, hereafter named *D4Z4-T* (D4Z4-Transcript), which arise from each D4Z4 repeat, and *DUX4FL* (DUX4 Full Length) pLAM-containing transcripts, the FSHD disease-associated mRNAs derived from the most telomere-proximal D4Z4 repeat (Fig. 1A and Supplemental_Fig_S1).

Our qPCR analyses (Fig 1C-D) showed that 4q35 genes are differentially expressed depending on their chromosomal position, confirming previous evidence [27,28,41,47]. Specifically, in control cells the centromere-proximal *ANT1*, *FAT1* and *FRG1* genes displayed high levels of expression, whereas *FRG2, DBE-T* and D4Z4-T were barely detectable. Consistent with the expected derepression of the locus associated with reduced D4Z4 copy number, *FRG2*, *DBE-T* and *D4Z4-T* transcripts were significantly upregulated in FSHD1 cells (Figure 1C and D). In contrast, *ANT1* and *FRG1* transcripts were found at comparable levels both in control and FSHD cells. Therefore, the loss of D4Z4 satellites in the FSHD patients’ cells correlated with an altered regulation of the telomere-proximal transcripts.

We could not reliably quantify the DUX4FL transcript via qPCR with commonly used primers [48,49] due to the very low amount of the detected amplicon (Ct values over 35) and because of the presence of multiple peaks in the melting curve analysis of the PCR products (Supplemental_Fig_S1). These observations are consistent with previous detection of DUX4FL transcripts only in a small percentage of FSHD-derived myoblasts (1 per 1000 cells) [44] and in cell lines only by using multi-step nested PCR [44,49–51].Since our aim was to conduct unbiased analyses of the physiological levels of 4q35 transcripts to compare their expression and regulation, we avoided pre-amplification steps.

The distinct regulation of genes located at different distances from the D4Z4 array prompted us to investigate the chromatin features of the 4q35 region. Chromatin immuno-precipitation (ChIP) experiments were performed in primary HPMs and HTCs using antibodies raised against histones tail modifications associated with transcriptional regulation: AcH3, AcH4 and H3K4me3, H3K9me3 and H3K27me3 (Figures 1E-F, values in Supplemental_Table1). We detected three classes of modification patterns. First, in both primary cell types, active chromatin marks were found at the *ANT1* and *FAT1* promoters within the centromere-proximal region of 4q35 region, with enrichment of AcH3, AcH4, and H3K4me3, and low levels of H3K9me3 and H3K27me3. Second, at the *FRG1* and *FRG2* promoters, both repressing and activating marks were detected. This “bivalent” pattern of histone modification is characteristic of “poised” promoters that are inactive but able to respond to external stimuli [52–54]. Specifically, we detected the enriched levels of AcH4 and H3K9me3 at the *FRG1* promoter in both cell types, at comparable levels in control and DRA-bearing cells. At the *FRG2* promoter, we observed comparable levels of AcH4 in control and FSHD-derived cells, whereas H3K9me3 enrichment was significatively more elevated in FSHD-derived myoblasts compared to control cells. A poised chromatin modification pattern at the *FRG2* promoter is a conserved feature in multiple cell types, as confirmed by Chromatin State Segmentation by HMM from studies by ENCODE/Broad [55] (Supplemental_Fig_S2). The third modification pattern observed occurred at the D4Z4 repeats themselves and was dominated by strong enrichment of H3K9me3 associated with low levels of the other chromatin marks analyzed (Figure 1E-F). A similar signature was also found at the gene desert region LILA5 (Supplemental_Fig_S3), as expected in heterochromatic regions. Notably, all three classes of modification patterns were largely similar comparing samples from healthy individuals and from FSHD patients with D4Z4 deleted alleles (DRAs). However, H3K9me3 levels were significantly increased at the *FRG2* promoter and at *D4Z4* in primary cells carrying a DRA [13,38]. This might seem to conflict with our observation that both *FRG2* and *D4Z4-T* were derepressed in cells carrying a DRA (Fig 1B and C). However, these data are consistent with previous results cells from patients with ICF (Immunodeficiency, Centromeric region instability, Facial anomalies syndrome), in which D4Z4 transcription is detected in spite of retention of H3K9me3 [56]. In sum, 4q35 contains three subdomains with euchromatic, poised, and heterochromatic features, arrayed in a centromere-to-telomere order.

### The *FRG2* promoter is activated by inhibition of histone acetylation or PARP1, in a manner regulated by D4Z4 repeat length

The detection of *FRG2* and D4Z4 transcription despite the enrichment of heterochromatin-associated histone marks at their promoters suggested complex modes of regulation. To investigate the role of chromatin modification across 4q35, we pharmacologically inhibited different classes of chromatin-modifying enzymes both in HTCs and HPMs (Fig. 2) and measured the RNA levels by RT-qPCR. We treated cells with trichostatin A (TSA), an inhibitor of class I, II, IV, histone deacetylases (HDACs) [57], nicotinamide (NAM) [58], an inhibitor of class III HDACs (sirtuins), or PJ34, an inhibitor of Poly (ADP-ribose) polymerase-1 (PARP-1) [59–61]. These treatments were performed either in presence or absence of 5-Aza-dC (Aza), an inhibitor of DNA methylation [57]. After these treatments, minor changes in expression of *ANT1*, *FAT1* and *FRG1* (Figure 2A-C and G-I) were observed. In contrast, strong transcriptional induction of *FRG2* was observed upon TSA and PJ34 treatments (Figure 2D and J, note the y-axis scale). The effects of both these compounds were enhanced by 5-aza-dC, indicating that DNA methylation contributes to *FRG2* silencing. Like the centromere-proximal genes, transcription of *DBE* and *D4Z4-T* transcripts was not induced by the selected compounds (Figure 2E-F and K-L). We conclude that the *FRG2* promoter is particularly sensitive to local chromatin modifications.

**Figure 2.**
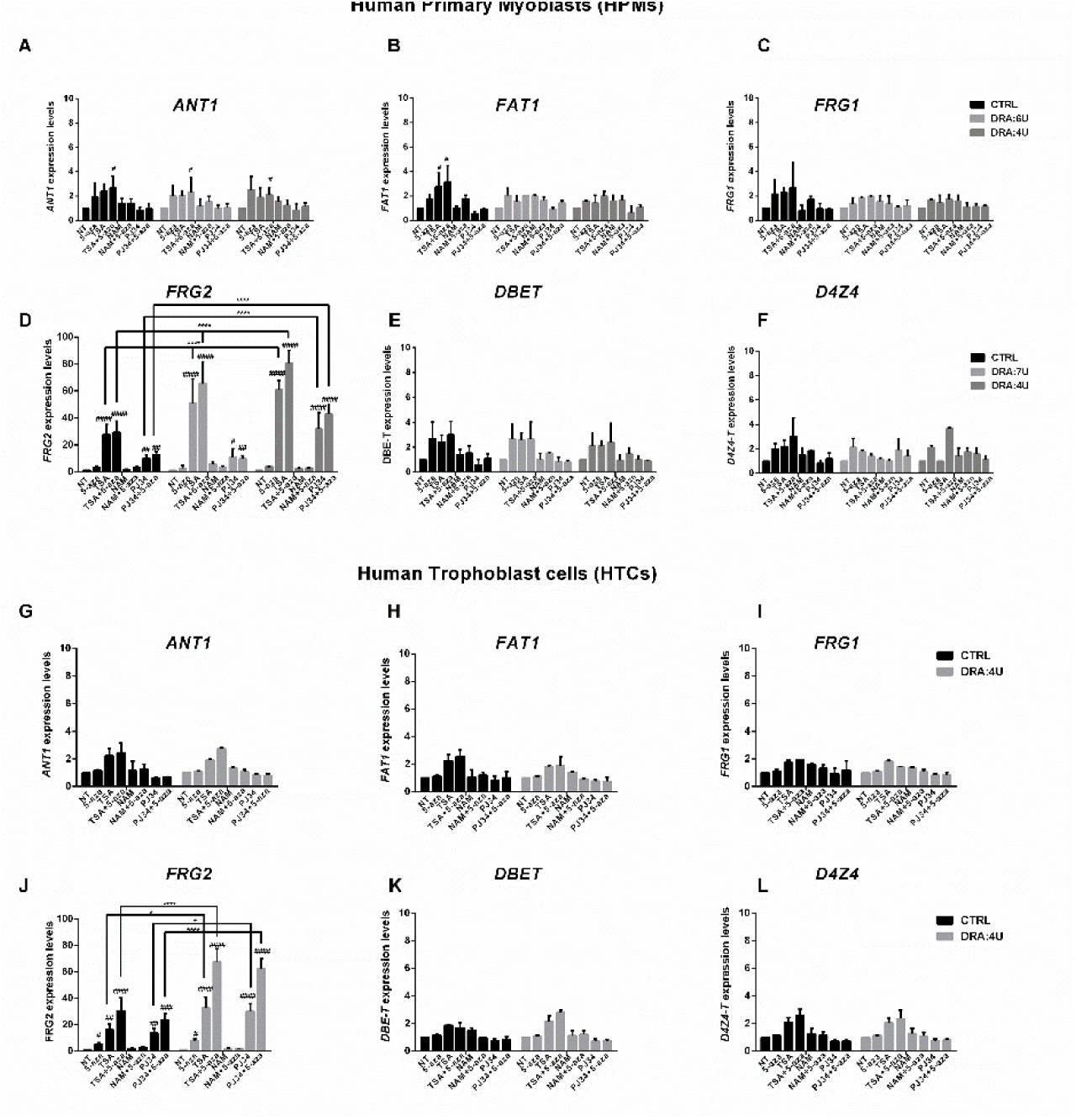
4q35 gene expression is affected by epigenetic drugs depending on 4q allele size. Expression data of 4q35 genes in human primary myoblasts (HPMs) (A-F) and in human trophoblasts cells (HTCs) (G-L) carrying a normal sized allele (CTRL (>10U)) or D4Z4 reduced alleles (DRA:7U and DRA:4U). Cells were treated or not treated with the indicated compounds: 5-Aza-2’-deoxycytidine (5-Aza-dC), Trichostatin A (TSA), nicotinamide (NAM), PARP inhibitor (PJ34). *ANT1* (A), *FAT1* (B), *FRG1* (C), *FRG2* (D), *DBE-T* (E) and *D4Z4-T* (F) RNAs were measured by RT q-PCR and normalized over the *RPLP0* reference gene. Experiments were done in triplicate and the results were analyzed using two-way Anova tests to perform multiple comparisons. Hashtags (#) indicate the statistical significance of data from treated samples compared to untreated samples (NT) in each group. Asterisks (*)indicate statistical significance of data from treated cells carrying DRA compared to the same treatment in control cells. P-value ranges are as follows: *, # 0.05<p-value<0.01; **, ## 0.01<p-value<0.001; ***, ### 0.001<p-value<0.0001; ****, #### p-value<0.0001.

To determine how TSA-induced transcriptional changes correlated with altered histone modifications, we performed ChIP experiments (Supplemental_Fig_S4, data in Supplemental Table 3). Both in control and DRA-bearing cells, TSA led to a general increase of ‘open chromatin’ marks (acetylated H3/H4 and H3K4me3) at 4q35 genes. In particular, and consistent with its transcriptional upregulation, we observed increased H3 and H4 acetylation at the poised *FRG2* promoter in HPMs and HTCs bearing one DRA allele (Supplemental_Fig_S4B-C and E). Together, our data indicate that histone acetylation and D4Z4 repeat length both contribute to the robust inducibility of the *FRG2* promoter.

### Transcription of *FRG2* and D4Z4 macrosatellite sequences is induced by genotoxic agents

To test whether 4q35 transcription was affected by a wider array of environmental perturbations, we analyzed the effects of genotoxic agents. We treated primary cells with Cisplatin (CIS) [62], Etoposide (ETO) [63] and Doxorubicin (DOXO) [64,65] (Figures 3 and Supplemental_FigS5). The centromere-proximal genes (*ANT1, FAT1 and FRG1*) were mildly repressed upon genotoxic injury. In contrast, expression of *FRG2, DUX4-T* and *DBE-T* increased significantly in the presence of all these compounds both in control and FSHD-derived cells. Additionally, genotoxic agents increased the amounts of 4q35 telomeric transcripts *FRG2*, *DBE-T* and *D4Z4-T* significantly more in HPMs and HTCs bearing a DRA in comparison with cells bearing normal sized D4Z4 alleles. We conclude that RNA levels from telomere-proximal 4q35 genes are induced by genotoxic agents, and the magnitude of this effect is increased by the presence of shortened D4Z4 arrays.

**Figure 3.**
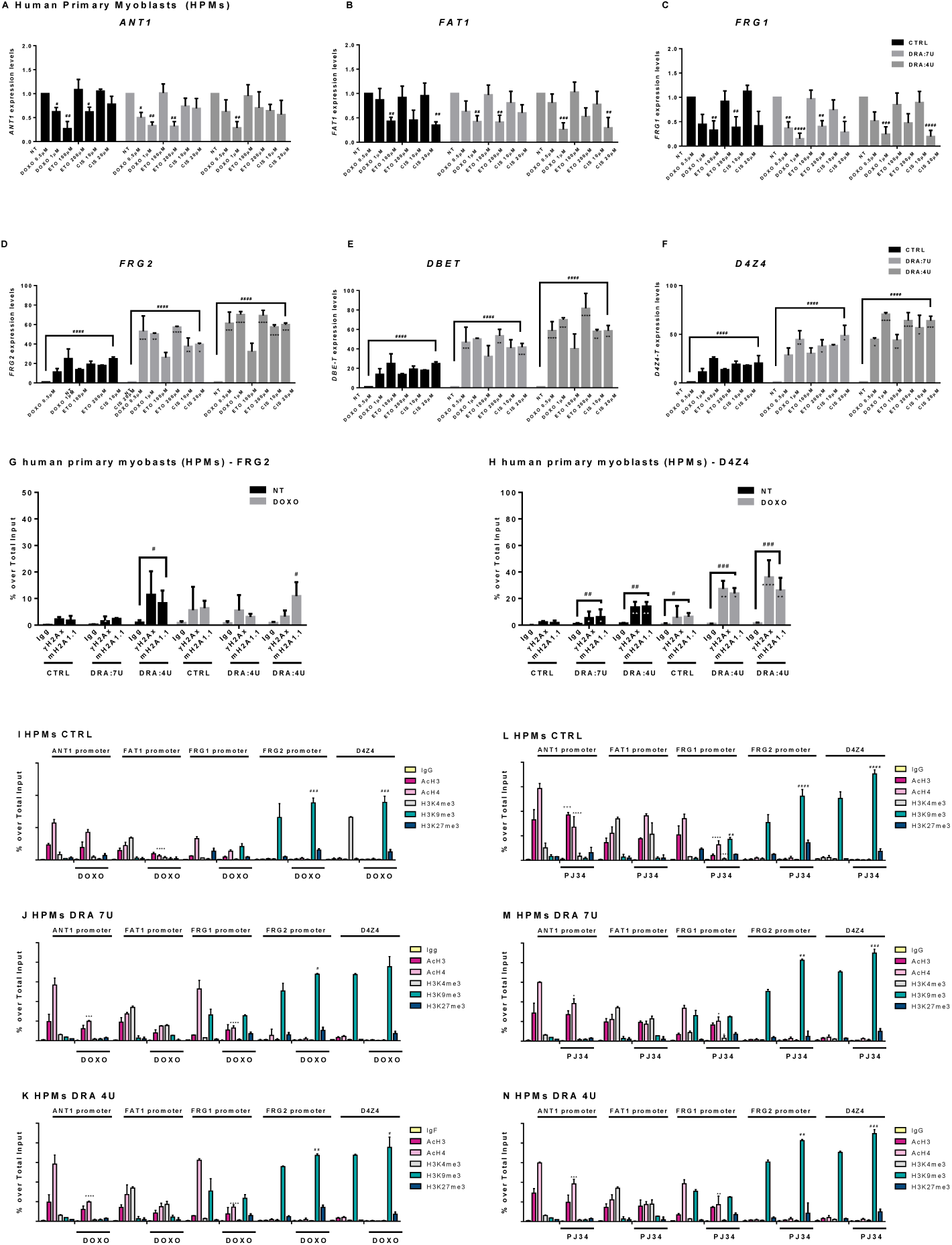
4q35 genes show different responsiveness to DNA damage depending on D4Z4 size and subtelomeric localization. Control HPMs and HPMs bearing 7U and 4U D4Z4 arrays were untreated or treated with genotoxic drugs: Doxorubicin (DOXO), Etoposide (ETO) and Cisplatin (CIS), at the reported concentrations. Expression data of *ANT1* (A), *FAT1* (B), *FRG1* (C), *FRG2* (D), *DBE-T* (E) and *D4Z4-T* (F) was evaluated 24h after treatments and normalized over *RPLP0* reference gene levels. Error bars represent standard deviation values for three independent replicates. Hashtags refer to statistical significance of treated samples in respect to not treated samples. Asterisks refer to statistical significance of treated cells carrying DRA in respect to the same treatment in control cells (NT) in each group. G-L) Chromatin immunoprecipitation assays (ChIP) conducted in control and DRA HPMs that were untreated or treated with Doxorubicin (G-I) or PJ34 (J-L). Antibodies directed to H3K4me3, H3K9me3, H3K27me3 and pan-acetylated Histone 3 and 4 (AcH3 and AcH4) were used, followed by qPCR amplification using primers described in Fig.1A. Anova statistical test with multiple comparison was performed (*0.05<p-value<0.01; ** 0.01<p-value<0.001; *** 0.001<p-value<0.0001; **** p-value<0.0001). Different symbols: * (asterisk) + sign (plus sign) and # (hashtag) refer to different antibodies used in ChIP experiments (*=AcH4; +=AcH34me3; #=H3K9me3 to show the statistical significance of data obtained in treated cells in respect to the same in not treated cells). M-N) Chromatin Immunoprecipitation (ChIP) in HPM cells that were untreated or treated with Doxorubicin (M) or PJ34 (N) . Antibodies directed to γH2Ax and macroH2A1.1 (mH2A1.1) were used followed by qPCR amplification of 4q35 genes as indicated. Anova statistical test with multiple comparison was performed (*0.05<p value<0.01; ** 0.01<p value<0.001; *** 0.001<p value<0.0001; **** P value<0.0001. * (asterisks) refer to each different antibody used in ChIP experiment to show the statistical significance of data obtained in treated cells in respect to the same in not treated cells.

To investigate more deeply the chromatin changes at 4q35 in response to DNA damage, we evaluated the amounts of histone isoforms that serve as DNA damage indicators: phosphorylated H2AX (γH2AX), which appears at DNA double-strand breaks, and macroH2A1.1, which is recruited to sites of DNA damage-induced PARP1 activation [65–67] (Figure 3G-H, values in Supplemental_Table2). At *FRG2*, we detected low levels of both these histones in the absence of DOXO treatment. At D4Z4, macroH2A.1 and 𝛄H2AX are present in basal levels untreated control cells, and these levels increased in cells carrying a 4U DRA or when DNA damage was induced by DOXO (Figure 3N). These observations suggest distinct chromatin architectures at 4q35 alleles that contain a DRA, in which the macrosatellite deletion renders the locus more accessible to DNA damaging agents and/or to DNA damage response factors.

We also evaluated histone modifications at the 4q35 genes in response to exogenous stresses (Figure 3I-N and Supplemental_Fig_S6, values in Supplemental_Table 2 and 4). Our analysis revealed that the DOXO-mediated increase of *FRG2, DBE-T* and *D4Z4-T* transcripts was not associated with local enrichment of histone acetylation or other transcription-associated histone modifications (Figure 3I-K and Supplemental Fig. S6A-B). Instead, a significant increase of H3K9me3 at *D4Z4* loci (p-values <0.001 and <0.01) was observed. Furthermore, H3K9me3 levels became significantly greater in cells bearing a DRA than in control cells, both at the *FRG2* promoter (p-value <0.001) and the *D4Z4* array (p-value <0.05). Treatment with PARP inhibitor PJ34 also induced a robust increase in H3me3K9 (p-value <0.001) at *FRG2* promoter and *D4Z4* in DRA cells (Figure 3L-N and Supplemental Fig S6C-D). Therefore, the increased RNA levels observed at the 4q35 telomere-proximal genes upon genotoxic stress or PARP inhibition are paradoxically accompanied by increased H3K9 methylation.

### Transcripts from telomere-proximal 4q35 genes are post-transcriptionally stabilized upon DNA damage

We observed that steady-state *FRG2* transcript levels were induced by TSA, in cells with DRA alleles this was accompanied by increased histone H3/H4 acetylation (Figure 2). In contrast, the increased transcript levels of *FRG2*, *DBE-T* and *D4Z4-T* upon genotoxic injury were not correlated with histone modifications typical for transcriptional activation increased H3K9me3 levels (Figure 3).

These observations were inconsistent with typical gene activation scenarios, but we reasoned that they could be consistent with post-transcriptional stabilization of telomeric 4q35 transcripts in presence of genotoxic damage. To test this, we treated control or FSHD HTCs with Actinomycin D (ActD), at concentration sufficient to inhibit transcription by both RNA polymerase I and II [68] and then evaluated the stability of 4q35 transcripts over time. Experiments were performed with ActD alone or in presence of DOXO (Figure 4). Notably, the expected increase in *FRG2, DBE-T* and *DUX4-T* transcript levels in the presence of DOXO was also observed when transcription was inhibited by ActD treatment (Fig. 4D-F), supporting our idea about post-transcriptional stabilization. Also, quantification of the data detected longer half-lives for these three telomere-proximal RNA species in cells with DRA alleles (Fig. 4A). We conclude that the major regulatory event for the 4q35 telomere-proximal transcripts upon genotoxic stress is post-transcriptional stabilization.

**Figure 4.**
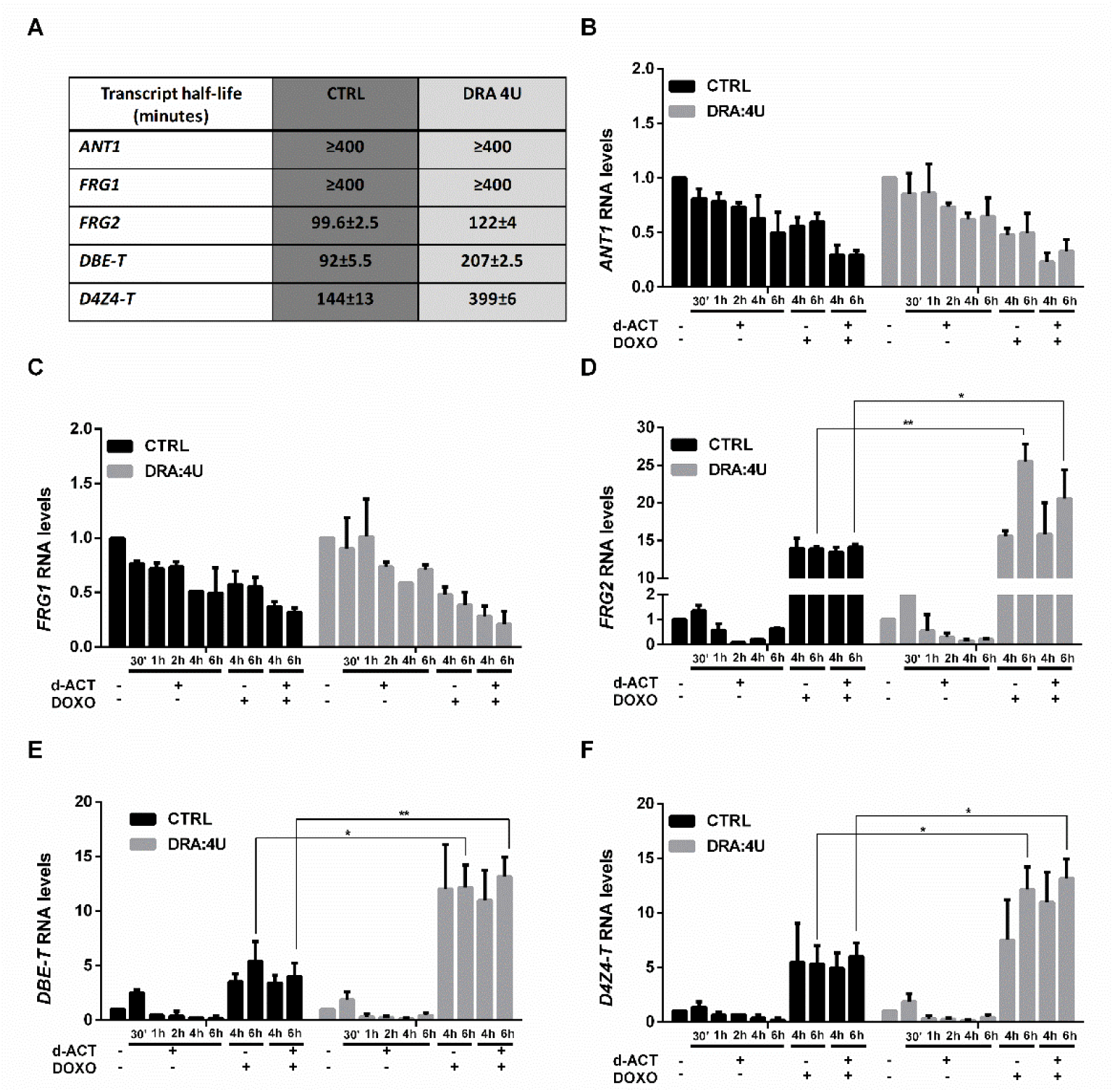
4q35 telomeric transcripts are stabilized upon DNA damage and transcriptional inhibition dependently on D4Z4 size reduction. A) Table reports the half-life of 4q35 transcripts measured after Actinomycin D (ActD) treatment in Control (CTRL) and DRA-containing (4U) HPMs. B-F) Cells were treated with ActD for the indicated times (30’, 1h, 2h, 4h and 6 h) in presence or absence of DOXO, and the levels of 4q35 gene transcripts were evaluated by qPCR. The half-lifes of each RNA was calculated as the time needed to reduce the transcript level to half (50%) of its initial abundance at time 0. Data shown are means ± s.e.m. of 3 replicates.

### Transcripts from 4q35 telomere-proximal genes are chromatin-associated

The observation that *FRG2*, *DBE-T* and *DUX4-T* transcript levels are affected by the same stimuli raises the question whether these RNAs have additional commonalities. Since repetitive element RNAs often function as components of chromatin fibers [69], we performed RNA fractionation experiments in primary control or FSHD-derived myoblasts (Fig. 5A-B). In both cell samples, *FRG2 and D4Z4-T* RNAs were enriched in the chromatin-associated fraction and behaved similarly to the previously characterized chromatin-associated transcripts lncDBE-T and TERRA [39,70]. As controls for the fractionation, we confirmed that the lncRNA NEAT1, was prevalently found in the nuclear fraction, and the protein-coding mRNA *GAPDH* was preferentially enriched in the cytoplasm.

**Figure 5.**
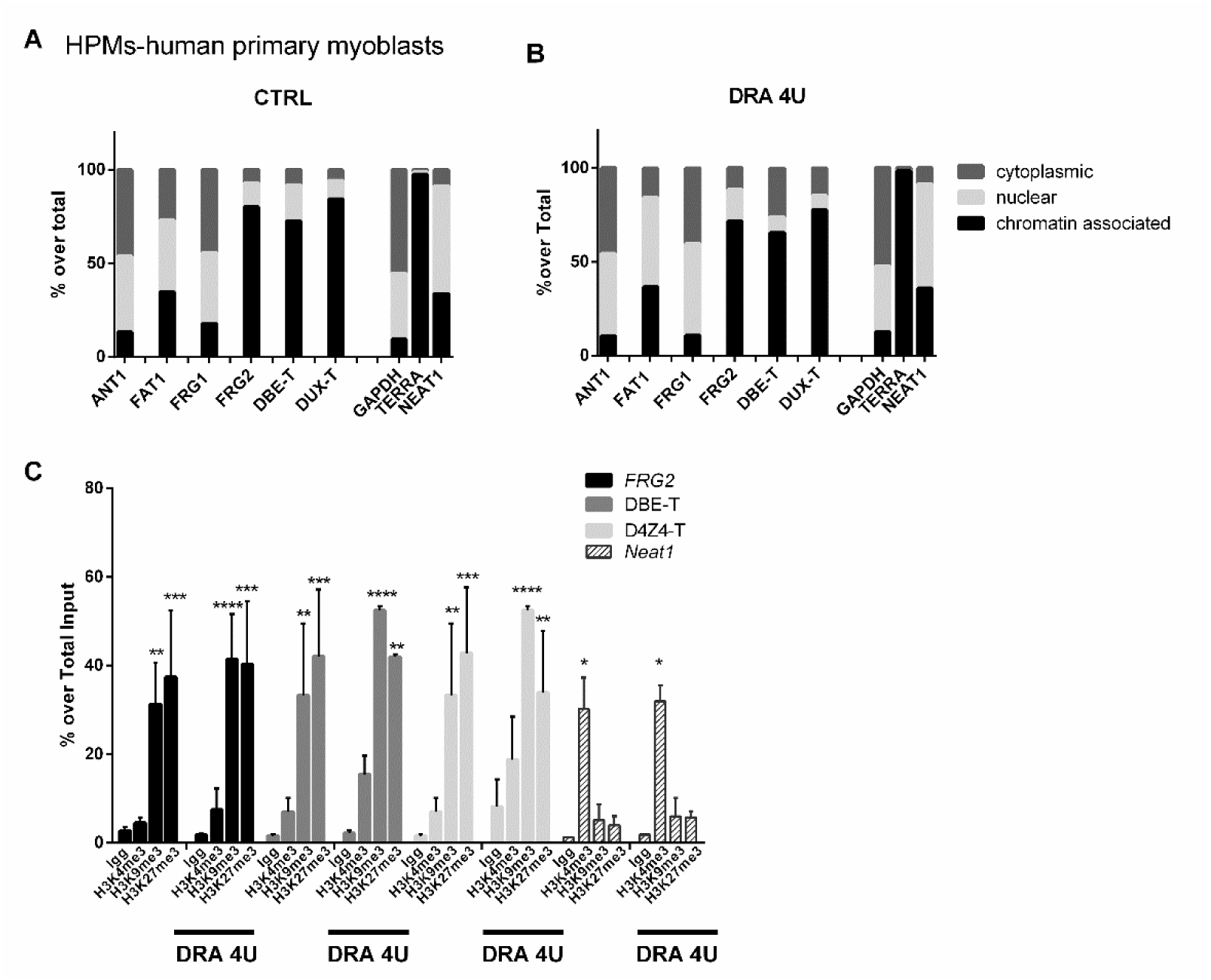
4q35 genes regulation upon different stimuli reflects architectural and epigentic patterns. A-B) RNA fractionation experiments were conducted in CTRL (A) and DRA (B) human primary myoblasts (HPMs). Transcripts from the indicated 4q35 genes were measured by qPCR analysis of cytoplasmic, nuclear and chromatin-associated RNA fractions, and the percentage detected in each fraction over total RNA was graphed. *GAPDH, TERRA* and *NEAT1* transcripts were also assessed as positive controls that are most enriched in cytoplasmic, chromatinic and nuclear fractions, respectively. C) ChRIP experiment performed in HPM cells. Antibodies directed to H3K4me3, H3K9me3, H3K27me3 were used to precipitate RNA from control and DRA cells. Data shown are means ± s.e.m. of 3 replicates. * (asterisks) refer to each different antibody used in ChRIP experiment to show the statistical significance of data obtained for each antibody over control IgG.

The chromatin association of the *FRG2*, *DBE-T* and *DUX4* transcripts was confirmed by Chromatin-RNA Immuno-Precipitation (ChRIP) [71] conducted in in primary myoblasts from control and FSHD subjects using H3K4me3, H3K9me3 and H3K27me3-specific antibodies (Fig. 5C). *FRG2*, *DBE-T* and *DUX4* transcripts were selectively and significatively enriched in H3K9me3 and H3K27me3-marked chromatin. As a control for the selectivity of our analysis, we confirmed that lncRNA Neat1, known to be associated with actively transcribed genes, was enriched in H3K4me3-marked but not H3K9me3 or H3K27me3-marked chromatin [72].

Together, our findings reveal that the telomere-proximal 4q35 genes share important regulatory features: their transcript levels are induced by genotoxic stress via post-transcriptional stabilization, and these RNAs are all chromatin-associated (Figure 6). Furthermore, the regulation of these telomeric transcripts is affected by the size of the D4Z4 sub-telomeric array (Figures 2 and 3). Therefore, the regulatory potentialc of this locus is expected to be variable in the human population.

**Figure 6.**
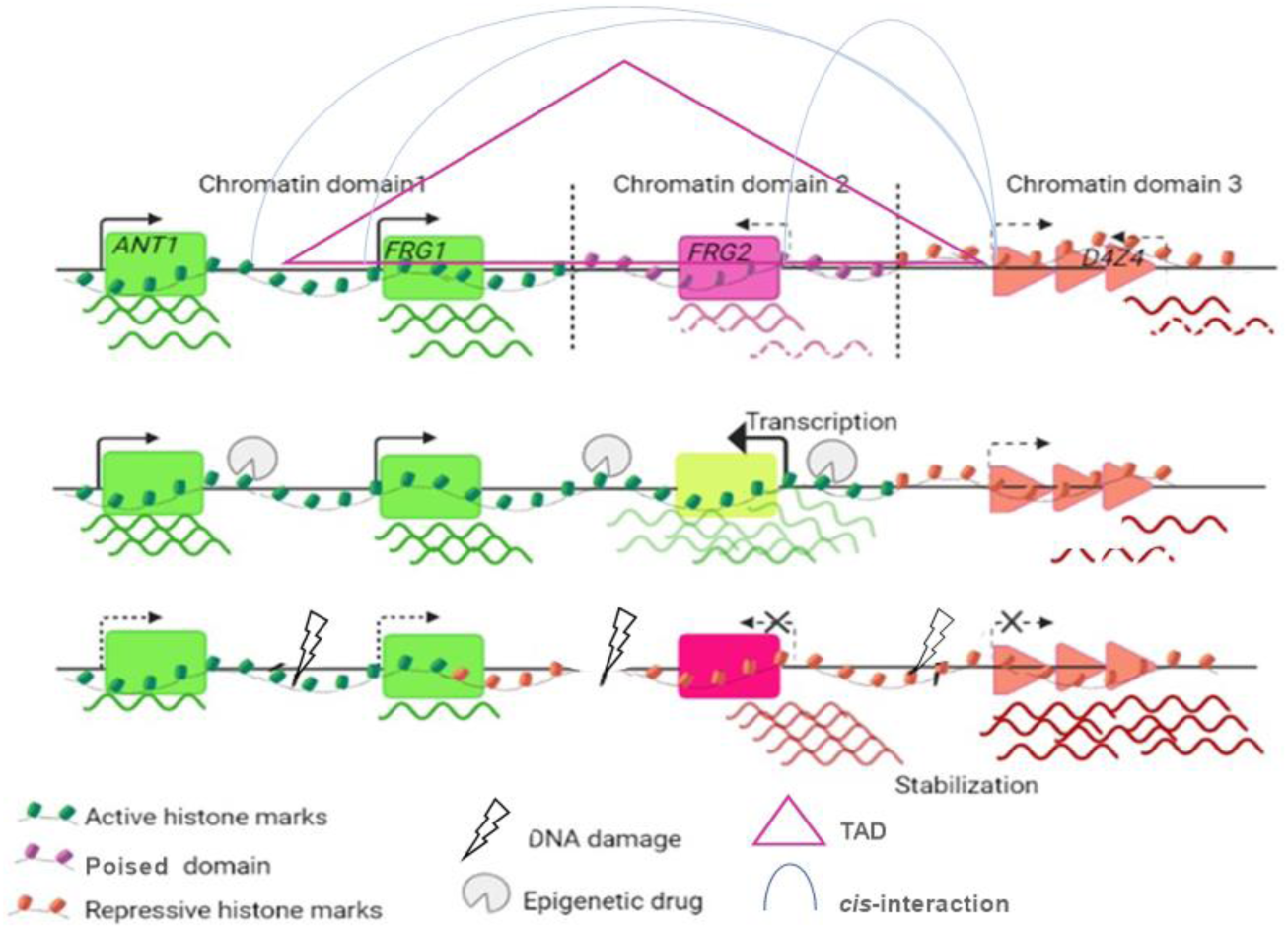
4q35 genes regulation upon different stimuli refects architectural and epigenetic patterns. Top diagram: A topological domain (TAD, indicated by the magenta triangle) at 4q35 includes the *FRG1* and *FRG2* genes [73](190–191 Mb of Chr 4). Additional *cis*-interactions between D4Z4 and nearby genes have also been reported [27,74] (curved lines). Our present study indicates functional subdomains within the 4q35 subtelomere, arrayed in a gradient along the chromosome. The different chromatin configurations at each subdomain correlate with the different response of these regions to external stimuli. The centromere-proximal genes *ANT1* and *FRG1* display active histone marks and are constitutively expressed at high levels (Chromatin domain 1). In contrast, the *FRG2* promoter displays a poised promoter (Chromatin domain 2). Finally, the telomeric genes at the *D4Z4* repeats (Chromatin domain 3) display repressive chromatin marks and are transcriptionally repressed in normal individuals in the absence of genotoxic stress. **Middle diagram:** Drug-induced epigenetic derepression (i.e. TSA treatment) results in enrichment of active histone marks at chromatin domain 1 promoters and a switch toward active chromatin at the FRG2 promoter leading to increased RNA levels. **Bottom diagram:** DNA damage (i.e. DOXO treatment), globally reduces the transcriptional activity across 4q35 and mediates a switch towards increased repressive chromatin markings at D4Z4 and the *FRG2* promoter. Additionally, transcripts from Chromatin domains 2 and 3 are stabilized through a posttranscriptional event. This model applies to control and to cells carrying a reduced D4Z4 allele, but the transcriptional or post transcriptional induction/stabilization rate is inversely correlated with D4Z4 size.

## DISCUSSION

### Novel modes of regulation of telomeric 4q35 transcripts provide insights for understanding clinical variability and low penetrance in FSHD

FSHD is a frequent myopathy, which has an estimated prevalence of 1 in 20,000 individuals [75]. Connecting the reduction of D4Z4 repeats with the development of FSHD is the major hurdle in understanding the molecular mechanism leading to disease. Inappropriate gene activation is considered the link between reduced copy number of the D4Z4 macrosatellite at 4q35 and the onset of FSHD, but clinical [76,77] and genetic [78] observations have challenged our understanding of the disease in finding a unifying model that fully addresses FSHD complexity. This complexity has been measured by stratification of heterozygous carriers of D4Z4 reduced alleles (DRAs) using a Comprehensive Clinical Evaluation Form (CCEF) [46,79]. This evaluation separates individuals into 4 categories following a straight-forward evaluation, from full penetrance to asymptomatic and atypical presentation, demonstrating a wide range of clinical phenotypes from people carrying similarly-sized DRAs. This phenotypic heterogeneity is also observed within families suggesting non-Mendelian factors may contribute [80–91]. Indeed, these phenotypic categorization are heterogeneous between probands and their first-degree relatives, with 50-75% of relatives remaining asymptomatic depending on the degree of kinship [80]. Interestingly in 35% of families in which a DRA with 7-8 repeat segregates only one affected individual (ie, the proband), is affected by disease, whereas the others are asymptomatic carriers; this finding holds regardless of the proband’s phenotypic category [92]. Together, these clinical datasets demonstrate highly variable penetrance of FSHD, and suggests that complex disease cofactors yet to be identified.

In the heterozygous state, a D4Z4 reduction might produce a subclinical sensitized condition that requires other epigenetic mechanisms or a contributing factor to cause overt myopathy. In some cases, it might be by the simultaneous heterozygosity for a different and recessive myopathy, as suggested by many reports in which the FSHD contractions are found in association with a second molecular defect [75,93–102]. Alternatively, as our findings are suggesting, environmental changes that affect chromatin modifications at 4q35 could generate an abnormal quantity of subtelomeric transcripts in cells with a DRA.

In this respect, the chromatin changes at 4q35 in cells bearing a DRA have been investigated to test for FSHD-specific D4Z4 *cis* and *trans* interactions [27,73,74] (Figure 6). For example, the 4q35 D4Z4 array possesses a CTCF-associated insulator which defines the boundaries of a D4Z4-proximal TAD [103]. (Figure 6). The D4Z4 array at 4q35 also tethers multiple telomeres to the nuclear envelope, inducing transcriptional repression of trans-associated genomic regions [25,103–105]. In this manner, a normal-sized D4Z4 array can contact several regions in a peripheral nuclear domain, possibly influencing transcription at multiple genomic regions. In contrast, in FSHD patients the shortened array displays reduced trans-interactions, with consequent transcriptional up-regulation of the distant loci [28]. Regarding regulation in cis, previous studies indicate that 4q telomere length regulates 4q35 genes as far as 5-Mb upstream of the *D4Z4* repeats, thus reinforcing the notion of a cooperative effect between telomere length and size of the *D4Z4* array [47] In sum, telomeric 4q35 genes are regulated by TAD boundaries and the length of both the D4Z4 array and telomere repeats [5,6].

In this study, we report additional levels of regulation of 4q35 gene expression. We found that the telomere-proximal *FRG2*, *DBE-T* and *D4Z4-T* genes are robustly sensitive to induction by genotoxic stress. Further, we discovered that transcripts from these genes are chiefly regulated by post-transcriptional stabilization, and that these regulations are affected by D4Z4-array size. We hypothesize that that 4q35 represents a case where subtelomere/telomere dynamics and their role in chromatin structure can influence the onset of pathologies and variability in clinical manifestations.

### D4Z4-derived transcripts in a genomic perspective

The precise basis by which over-expression of 4q35 genes results in FSHD remains to be determined. Here we report that D4Z4-derived transcripts are stabilized upon genotoxic stress, revealing an additional level of regulation involving post-transcriptional events. Consistent with this discovery are earlier hypotheses that Nonsense Mediated Decay (NMD) acts as an endogenous suppressor of DUX4FL mRNA [106], or that NMD is globally impaired in FSHD cells [107]. We note that persistent DNA damage inhibits NMD through p38α MAPK pathway activation [108], which would further link our data to the NMD hypotheses. Therefore, more investigation is required to characterize how NMD regulation by DNA damage affects RNA targets in cell-specific contexts, particularly regarding FSHD phenotypes [109].

Our data confirm that D4Z4-derived RNAs display elevated levels and are stabilized in FSHD cells, raising questions about their roles during DNA damage. As components of the chromatin fiber, these RNAs could favor regulatory *cis* chromatin interactions at 4q35 [39], as well as *trans* interactions with other repetitive elements interspersed across the human genome. Furthermore, as D4Z4-like sequences are highly polymorphic and account for several hundreds of Kbs in each individual genome [16], further studies will be required to determine the extent of D4Z4 and D4Z4-like RNA molecules produced, how these differ in different individuals, and how their interaction patterns impact genome function. Therefore, FSHD offers a valuable natural model to understand the complex interplay between tandemly arrayed repats and their function in genomic architecture and phenotypic heterogeneity. The development of long read DNA and RNA sequencing technologies offers unprecedented possibilities for in-depth molecular phenotyping and for the interpretation of results in a multidimensional perspective.

### A role for DNA repetitive elements in personalized medicine

One of the current challenges in molecular medicine is to understand how DNA variations in non-coding sequences translate into phenotypic variability among individuals. Repetitive DNA elements represent 56% to 69% of the human genome [1,2,14,110,111]. Although macrosatellite repeats have been less well studied than many repeat classes, there is increasing evidence for a strong correlation between macrosatellite copy number, epigenetic modifications and local gene expression (references). Thus, macrosatellites provide an example of repeat-induced gene silencing as a mechanism of gene regulation in humans [112–114].

Our work demonstrates that the D4Z4 macrosatellite array alters transcriptional responses to drugs at nearby genes, and that the number of repetitive elements modulates these changes. It is thus possible that what we observe at the 4q35 D4Z4 locus may occur at other macrosatellites interspersed within the genome. These sequences are highly polymorphic between individuals and heritable. Therefore, correlations between macrosatellite composition, size and their responsiveness to drugs could facilitate the understanding of how a person’s repetitive DNA profile contributes to disease susceptibility and could increase our ability to predict the results of specific medical treatments. In this manner, better knowledge of the biological roles of repeats may offer substantial tools for personalized medicine.

## MATERIAL AND METHODS

### Cell Culture and Drugs Treatment

Primary trophoblasts cell culture were established immediately and after Chorionic villus sampling (HTCs), and cells were grown in Ham’s F10 medium (Corning) plus L-glutamine, 20% FBS, and 1% Penicillin/Streptomycin. Healthy and FSHD-derived human primary myoblasts (HPMs) were cultured in DMEM, added with 20% FBS, 1% L-glutamine, 1% Penicillin/Streptomycin, 2 ng/mL epidermal growth factor (EGF) and 25 ng/mL of fibroblast growth factor (FGF). Both cell lines and primary cells were grown on T75 flasks in a humidified atmosphere at 37 °C with 6% CO_2_. Cells were treated with TSA (Sigma;T8552), PJ34 (Sigma; 528150) %-aza-cytidine (Sigma; A3656); NAM (Sigma; N0636); Doxorubicin (Sigma; D1515); Actinomycin D 1 µg/ml (Sigma; SBR00013).

### RNA extraction and real time quantitative RT-PCR (qRT-PCR)

Total cellular RNA was obtained using a PureLink RNA Mini Kit (Thermo Fisher Scientifc cat #12183018A) according to the manufacturer’s instructions. DNAse digestion and cDNA synthesis was performed by using Maxima H-cDNA Synthesis Master Mix, with dsDNase (Thermo Fisher M1682). Specific mRNA expression was assessed by qRT-PCR (iTaq Universal SYBR® Green Supermix, BIORAD#1725120 in a CFX connect Real Time Machine BIORAD) using primers listed in Supplemental Table M1, normalized over RPLP0 housekeeping mRNA.

### Statistical analyses

All data presented in figures were performed at least in triplicate and expressed as means ±SEM When making multiple statistical comparisons, one-way ANOVA with Tukey or Dunnett’s post-hoc tests was used for normally distributed data. All analyses were conducted using Prism software (GraphPad Software Inc.).

### ChIP and real time quantitative RT-PCR (qRT-PCR)

Chromatin immunoprecipitation (ChIP) assays were performed both in cell lines and in primary cells as described earlier [115] using specific antibodies as listed in Supplemental Table M2. Immunoprecipitates from at least three independent cell samples were analyzed by quantitative real-time PCR (qPCR) as described above. Enrichment of amplified DNA sequences (primers listed in Supplemental Table M1) in immunoprecipitates was calculated as the ratio between the DNA amount in immunoprecipitation samples and that in the total input chromatin.

### RNA fractionation

Control and FSHD derived myoblasts cell pellets (1 million of cells) were lysed with 175 μl of cold cytoRNA solution (50 mM Tris HCl pH 8.0; 140 mM NaCl; 1.5 mM MgCl2; 0,5% NP-40; 2mM Vanadyl Ribonucleoside Complex; Sigma) and incubated 5′ in ice. Cell suspension was centrifuged at 4°C and 300 g for 2′ and the supernatant, corresponding to the cytoplasmic fraction, was transferred into a new tube and stored in ice. The pellet containing nuclei were extracted with 175μl of cold nucRNA solution (50 mM Tris HCl pH 8.0; 500 mM NaCl; 1.5 mM MgCl2; 0,5% NP-40; 2mM Vanadyl Ribonucleoside Complex) and incubated 5′ on ice. The suspension was centrifuged at 4°C and 16360 g for 2′ and the supernatant, corresponding to the nuclear-soluble fraction, was transferred into a new tube and stored in ice. The remaining pellet was collected as the chromatin-associated fraction.Total RNA from the cytoplasmic and nuclear fractions was extracted by using PureLink RNA MiniKit (Invitrogen) following the manufacturer’s instructions for the RNA extraction from aqueous solutions. The pellet containing the chromatin-associated fraction was extracted with the standard procedure described above for RNA extraction.

### Chromatin-RNA Precipitation (ChRIP)

Chromatin RNA immunoprecipitation (ChRIP) was performed as previously described [71] using anti-H3K4me3, H3K9me3 H3K27me3 antibodies as reported in TableM2. 3 × 10^6^ HPM cells were used for each IP). RNA was extracted and qPCR performed as described above. Ten percent input was used to calculate the percentage of transcript bound to chromatin compared to the negative control IgG.

## Author Contributions

VS, MS contributed to study design, molecular analysis, data collection, data analysis and interpretation, literature search, preparing of figures/tables and manuscript writing. FL performed molecular biology experiments. PDK contributed to data interpretation and manuscript editing. RT contributed to conception and study design, data interpretation, literature search and manuscript writing.

## Supporting information

Salsi et al Supplemental Tables

Salsi et al Supplemental Figures and legends

## Acknowledgements

We are indebted to all patients and their families for participating in this study. We wish to express our sincere gratitude and recognition to Professor Michael R. Green, who sadly left us too soon for his significant contributions to our research.

## Funding

National Institutes of Health, (Ro1ns0475840)(to RT), FSHD Global Research Foundation (to RT), and NIH U01 CA260669 (to PDK).

## Conflicts of Interest

The authors declare no conflict of interest.

## SUPPLEMENTAL FIGURES

SF1 Expression analysis of DUX4-T and DUX4FL. A) Schematic representation of the last D4Z4 unit, the adjacent pLAM region and the distal exons. The DUX4 ORF is contained in the first exon in each repeat. The pLAM sequence is only present in the DUX4-FL transcript. Primers used in for qPCR are indicated in different colors. B-D) qPCR reports of DUX-T and DUX-FL amplification by using the indicated primer pairs, melting cure graphs are visible in the upper part of each graph, melting curve and quantification cycle (C_q_) reports are indicated in the table below for human primary myoblasts (HPMs) and human primary trophoblast cells (here HPFs). For DUX-FL we used two different primer sets (pLAM1, purple and pLAM2 blue) used in previous works [48,49].

SF2 Chromatin State Segmentation by HMM from ENCODE/Broad; Hg19chr4:190,793,373-191,038,972.

SF3 Epigenetic status of 4q35-associated gene desert region. Chromatin immunoprecipitation assays (ChIP) conducted in (A) HPMs and HTCs (B) carrying a normal sized (control) and reduced (DRA) D4Z4 alleles. Antibodies directed to H3K4me3, H3K9me3, H3K27me3 and pan-acetylated Histone 3 and 4 (AcH3 and AcH4) were used, followed by qPCR amplification using primers described in Fig1.A.

SF4. G-H) Chromatin immunoprecipitation assays (ChIP) conducted in HPMs (A-C) and HTCs (D-E) carrying a normal sized (control) and reduced (DRA) D4Z4 alleles (B,C,E) and treated or not with TSA. Antibodies directed to H3K4me3, H3K9me3, H3K27me3 and pan-acetylated Histone 3 and 4 (AcH3 and AcH4) were used, followed by qPCR amplification using primers described in Fig1.A. Anova statistical test with multiple comparison was performed (*0.05<p value<0.01; ** 0.01<p value<0.001; *** 0.001<p value<0.0001; **** P value<0.0001): $ (dollar symbol), and * (asterisk). Dollar symbols and asterisks indicate the statistical significance of data obtained in TSA treated cells in respect to the same antibody enrichment in not treated cells. 4q35 regions amplified in qPCR following ChIP are indicated in the upper part of each graph.

SF5 Control HTCs and HTCs bearing 6U D4Z4 array were untreated or treated with genotoxic drugs: Doxorubicin (DOXO), Etoposide (ETO) and Cisplatin (CIS), at the reported concentrations. Expression data of *ANT1* (A), *FAT1* (B), *FRG1* (C), *FRG2* (D), *DBE-T* (E) and *D4Z4-T* (F) was evaluated 24h after treatments and normalized over *RPLP0* reference gene levels. Error bars represent standard deviation values for three independent replicates.

SF6 A-D)Chromatin immunoprecipitation assays (ChIP) conducted in control and DRA HTCs treated or not with Doxorubicin (A,B) or PJ34 (C,D). Antibodies directed to H3K4me3, H3K9me3, H3K27me3 and pan-acetylated Histone 3 and 4 (AcH3 and AcH4) were used, followed by qPCR amplification using primers described in Fig1.A. Anova statistical test with multiple comparison was performed (*0.05<p value<0.01; ** 0.01<p value<0.001; *** 0.001<p value<0.0001; **** P value<0.0001). Different symbols: * (asterisk) and # (hashtag) refer to different antibodies used in ChIP experiments (*=AcH4; #=H3K9me3 to show the statistical significance of data obtained in treated cells in respect to the same in not treated cells). . Error bars represent standard deviation values for three independent replicates.

## Notes

### Competing Interest Statement

The authors have declared no competing interest.

## REFERENCES

1. Biscotti MA, Olmo E, Heslop-Harrison JSP. Repetitive DNA in eukaryotic genomes. Chromosome Res. 2015;23:415–20.

2. Liao X, Zhu W, Zhou J, Li H, Xu X, Zhang B, et al. Repetitive DNA sequence detection and its role in the human genome. Commun Biol. 2023;6:954.

3. Cremer T, Cremer C. Chromosome territories, nuclear architecture and gene regulation in mammalian cells. Nat Rev Genet. 2001;2:292–301.

4. Schneider R, Grosschedl R. Dynamics and interplay of nuclear architecture, genome organization, and gene expression. Genes Dev. 2007;21:3027–43.

5. Robin JD, Ludlow AT, Batten K, Magdinier F, Stadler G, Wagner KR, et al. Telomere position effect: regulation of gene expression with progressive telomere shortening over long distances. Genes Dev. 2014;28:2464–76.

6. Stadler G, Rahimov F, King OD, Chen JCJ, Robin JD, Wagner KR, et al. Telomere position effect regulates DUX4 in human facioscapulohumeral muscular dystrophy. Nat Struct Mol Biol. 2013;20:671–8.

7. Misteli T. The long reach of telomeres. Genes Dev. 2014;28:2445–6.

8. Laberthonnière C, Magdinier F, Robin JD. Bring It to an End: Does Telomeres Size Matter? Cells. 2019;8:30.

9. Deutekom JCT van, Wljmenga C, Tlenhoven EAE van, Gruter AM, Hewitt JE, Padberg GW, et al. FSHD associated DNA rearrangements are due to deletions of integral copies of a 3.2 kb tandemly repeated unit. Human Molecular Genetics. 1993;2:2037–42.

10. Lunt PW, Jardine PE, Koch M, Maynard J, Osborn M, Williams M, et al. Phenotypic-genotypic correlation will assist genetic counseling in 4q35-facioscapulohumeral muscular dystrophy. Muscle & nerve Supplement. 1995;S103–9.

11. Chadwick BP. Macrosatellite epigenetics: The two faces of DXZ4 and D4Z4. Chromosoma. 2009;118:675–81.

12. Bakker E, Wijmenga C, Vossen RHAM, Padberg GW, Hewitt J, van Der Wielen M, et al. The FSHD-linked locus D4F104S1 (p13E-11) ON 4q35 has a homologue on 10qter. Muscle & Nerve. 1995;18:S39–44.

13. Zeng W, Chen YY, Newkirk DA, Wu B, Balog J, Kong X, et al. Genetic and Epigenetic Characteristics of FSHD-Associated 4q and 10q D4Z4 that are Distinct from Non-4q/10q D4Z4 Homologs. Human Mutation. 2014;35:998–1010.

14. Nurk S, Koren S, Rhie A, Rautiainen M, Bzikadze AV, Mikheenko A, et al. The complete sequence of a human genome. Science. 2022;376:44–53.

15. Liao W-W, Asri M, Ebler J, Doerr D, Haukness M, Hickey G, et al. A draft human pangenome reference. Nature. 2023;617:312–24.

16. Salsi V, Chiara M, Pini S, Kuś P, Ruggiero L, Bonanno S, et al. A human pan-genomic analysis reconfigures the genetic and epigenetic make up of facioscapulohumeral muscular dystrophy [Internet]. medRxiv; 2023 [cited 2023 Aug 17]. p. 2023.06.13.23291337.

17. Hewitt JE, Lyle R, Clark LN, Valleley EM, Wright TJ, Wijmenga C, et al. Analysis of the tandem repeat locus D4Z4 associated with facioscapulohumeral muscular dystropothhy. Human Molecular Genetics. 1994;3:1287–95.

18. Winokur ST, Bengtsson U, Feddersen J, Mathews KD, Weiffenbach B, Bailey H, et al. The DNA rearrangement associated with facioscapulohumeral muscular dystrophy involves a heterochromatin-associated repetitive element: Implications for a role of chromatin structure in the pathogenesis of the disease. Chromosome Research. 1994;2:225–34.

19. Winokur ST, Bengtsson U, Vargas JC, Wasmuth JJ, Altherr MR. The evolutionary distribution and structural organization of the homeobox-containing repeat D4Z4 indicates a functional role for the ancestral copy in the FSHD region. Human Molecular Genetics. 1996;5:1567–75.

20. Bizhanova A, Kaufman PD. Close to the edge: Heterochromatin at the nucleolar and nuclear peripheries. Biochimica et Biophysica Acta (BBA) - Gene Regulatory Mechanisms. 2021;1864:194666.

21. Santoro R, Grummt I. Molecular mechanisms mediating methylation-dependent silencing of ribosomal gene transcription. Molecular Cell. 2001;8:719–25.

22. Peng T, Hou Y, Meng H, Cao Y, Wang X, Jia L, et al. Mapping nucleolus-associated chromatin interactions using nucleolus Hi-C reveals pattern of heterochromatin interactions. [cited 2023 Apr 18].

23. Masny PS, Bengtsson U, Chung S-A, Martin JH, van Engelen B, van der Maarel SM, et al. Localization of 4q35.2 to the nuclear periphery: is FSHD a nuclear envelope disease? Hum Mol Genet. 2004;13:1857–71.

24. Tam R, Smith KP, Lawrence JB. The 4q subtelomere harboring the FSHD locus is specifically anchored with peripheral heterochromatin unlike most human telomeres. J Cell Biol. 2004;167:269–79.

25. Guelen L, Pagie L, Brasset E, Meuleman W, Faza MB, Talhout W, et al. Domain organization of human chromosomes revealed by mapping of nuclear lamina interactions. Nature. 2008;453:948–51.

26. Petrov A, Pirozhkova I, Carnac G, Laoudj D, Lipinski M, Vassetzky YS. Chromatin loop domain organization within the 4q35 locus in facioscapulohumeral dystropny patients versus normal human myoblasts. Proceedings of the National Academy of Sciences of the United States of America. 2006;103:6982–7.

27. Robin JD, Ludlow AT, Batten K, Gaillard M-C, Stadler G, Magdinier F, et al. SORBS2 transcription is activated by telomere position effect-over long distance upon telomere shortening in muscle cells from patients with facioscapulohumeral dystrophy. Genome Res. 2015;25:1781–90.

28. Cortesi A, Pesant M, Sinha S, Marasca F, Sala E, Gregoretti F, et al. 4q-D4Z4 chromatin architecture regulates the transcription of muscle atrophic genes in facioscapulohumeral muscular dystrophy. Genome Research. 2019;29:883–95.

29. Huichalaf C, Micheloni S, Ferri G, Caccia R, Gabellini D. DNA methylation analysis of the macrosatellite repeat associated with FSHD muscular dystrophy at single nucleotide level. PLoS ONE. 2014;9.

30. Salsi V, Magdinier F, Tupler R. Does DNA methylation matter in FSHD? Genes. 2020;11:258.

31. Nikolic A, Jones TI, Govi M, Mele F, Maranda L, Sera F, et al. Interpretation of the Epigenetic Signature of Facioscapulohumeral Muscular Dystrophy in Light of Genotype-Phenotype Studies. Int J Mol Sci. 2020;21:2635.

32. Van Overveld PGM, Enthoven L, Ricci E, Rossi M, Felicetti L, Jeanpierre M, et al. Variable hypomethylation of D4Z4 in facioscapulohumeral muscular dystrophy. Annals of Neurology. 2005;58:569–76.

33. Butterfield RJ, Dunn DM, Duvall B, Moldt S, Weiss RB. Deciphering D4Z4 CpG methylation gradients in fascioscapulohumeral muscular dystrophy using nanopore sequencing. bioRxiv. 2023;2023.02.17.528868.

34. Erdmann H, Scharf F, Gehling S, Benet-Pagès A, Jakubiczka S, Becker K, et al. Methylation of the 4q35 D4Z4 repeat defines disease status in facioscapulohumeral muscular dystrophy. Brain : a journal of neurology [Internet]. 2022 [cited 2023 Apr 2].

35. Gould T, Jones TI, Jones PL. Precise Epigenetic Analysis Using Targeted Bisulfite Genomic Sequencing Distinguishes FSHD1, FSHD2, and Healthy Subjects. Diagnostics (Basel, Switzerland) [Internet]. 2021 [cited 2023 Apr 2];11.

36. Das S, Chadwick BP. Influence of repressive histone and DNA methylation upon D4Z4 transcription in non-myogenic cells. Kyba M, editor. PLoS ONE. 2016;11:e0160022.

37. Vakoc CR, Mandat SA, Olenchock BA, Blobel GA. Histone H3 lysine 9 methylation and HP1γ are associated with transcription elongation through mammalian chromatin. Molecular Cell. 2005;19:381–91.

38. Zeng W, De Greef JC, Chen YY, Chien R, Kong X, Gregson HC, et al. Specific loss of histone H3 lysine 9 trimethylation and HP1γ/cohesin binding at D4Z4 repeats is associated with facioscapulohumeral dystrophy (FSHD). Ferguson-Smith AC, editor. PLoS Genetics. 2009;5:e1000559.

39. Cabianca DS, Casa V, Bodega B, Xynos A, Ginelli E, Tanaka Y, et al. A long ncRNA links copy number variation to a polycomb/trithorax epigenetic switch in fshd muscular dystrophy. Cell. 2012;149:819–31.

40. Campbell AE, Shadle SC, Jagannathan S, Lim J-W, Resnick R, Tawil R, et al. NuRD and CAF-1-mediated silencing of the D4Z4 array is modulated by DUX4-induced MBD3L proteins. eLife [Internet]. 2018 [cited 2024 Mar 8];7.

41. Gabellini D, Green MR, Tupler R. Inappropriate gene activation in FSHD: A repressor complex binds a chromosomal repeat deleted in dystrophic muscle. Cell. 2002;110:339–48.

42. Salsi V, Vattemi GNA, Tupler RG. The FSHD jigsaw: are we placing the tiles in the right position? Curr Opin Neurol. 2023;36:455–63.

43. Banerji CRS, Zammit PS. Pathomechanisms and biomarkers in facioscapulohumeral muscular dystrophy: roles of DUX4 and PAX7. EMBO Molecular Medicine. 2021;13.

44. Snider L, Geng LN, Lemmers RJLF, Kyba M, Ware CB, Nelson AM, et al. Facioscapulohumeral dystrophy: Incomplete suppression of a retrotransposed gene. Pearson CE, editor. PLoS Genetics. 2010;6:1–14.

45. Lemmers RJLF, Van Der Vliet PJ, Klooster R, Sacconi S, Camaño P, Dauwerse JG, et al. A unifying genetic model for facioscapulohumeral muscular dystrophy. Science. 2010;329:1650–3.

46. Bettio C, Salsi V, Orsini M, Calanchi E, Magnotta L, Gagliardelli L, et al. The Italian National Registry for FSHD: An Enhanced Data Integration and an Analytics Framework Towards Smart Health Care and Precision Medicine for a Rare Disease. 2021;

47. Robin JD, Ludlow AT, Batten K, Magdinier F, Stadler G, Wagner KR, et al. Telomere position effect: regulation of gene expression with progressive telomere shortening over long distances. Genes Dev. 2014;28:2464–76.

48. Lemmers RJLF, Van Der Vliet PJ, Blatnik A, Balog J, Zidar J, Henderson D, et al. Chromosome 10q-linked FSHD identifies DUX4 as principal disease gene. Journal of Medical Genetics. 2022;59:180–8.

49. Das S, Chadwick BP. Influence of repressive histone and DNA methylation upon D4Z4 transcription in non-myogenic cells. Kyba M, editor. PLoS ONE. 2016;11:e0160022.

50. Jones TI, Chen JCJ, Rahimov F, Homma S, Arashiro P, Beermann ML, et al. Facioscapulohumeral muscular dystrophy family studies of DUX4 expression: Evidence for disease modifiers and a quantitative model of pathogenesis. Human Molecular Genetics. 2012;21:4419–30.

51. Jones TI, King OD, Himeda CL, Homma S, Chen JCJ, Beermann ML, et al. Individual epigenetic status of the pathogenic D4Z4 macrosatellite correlates with disease in facioscapulohumeral muscular dystrophy. Clinical Epigenetics. 2015;7:37.

52. Crispatzu G, Rehimi R, Pachano T, Bleckwehl T, Cruz-Molina S, Xiao C, et al. The chromatin, topological and regulatory properties of pluripotency-associated poised enhancers are conserved in vivo. Nat Commun. 2021;12:4344.

53. Boltsis I, Grosveld F, Giraud G, Kolovos P. Chromatin Conformation in Development and Disease. Front Cell Dev Biol [Internet]. 2021 [cited 2024 Mar 12];9.

54. Becker JS, McCarthy RL, Sidoli S, Donahue G, Kaeding KE, He Z, et al. Genomic and Proteomic Resolution of Heterochromatin and Its Restriction of Alternate Fate Genes. Molecular Cell. 2017;68:1023–1037.e15.

55. Ernst J, Kheradpour P, Mikkelsen TS, Shoresh N, Ward LD, Epstein CB, et al. Mapping and analysis of chromatin state dynamics in nine human cell types. Nature. 2011;473:43–9.

56. Kondo T, Bobek MP, Kuick R, Lamb B, Zhu X, Narayan A, et al. Whole-genome methylation scan in ICF syndrome: Hypomethylation of non-satellite DNA repeats D4Z4 and NBL2. Human Molecular Genetics. 2000;9:597–604.

57. Yang F, Zhang L, Li J, Huang J, Wen R, Ma L, et al. Trichostatin A and 5-azacytidine both cause an increase in global histone H4 acetylation and a decrease in global DNA and H3K9 methylation during mitosis in maize. BMC plant biology. 2010;10:178.

58. Rymarchyk S, Kang W, Cen Y. Substrate-Dependent Sensitivity of SIRT1 to Nicotinamide Inhibition. Biomolecules. 2021;11:312.

59. Rouleau M, Patel A, Hendzel MJ, Kaufmann SH, Poirier GG. PARP inhibition: PARP1 and beyond. Nature Reviews Cancer. 2010;10:293–301.

60. Zha S, Wang F, Li Z, Ma Z, Yang L, Liu F. PJ34, a PARP1 inhibitor, promotes endothelial repair in a rabbit model of high fat diet-induced atherosclerosis. Cell Cycle. 2019;18:2099–109.

61. Ying W, Alano CC, Garnier P, Swanson RA. NAD+ as a metabolic link between DNA damage and cell death. Journal of Neuroscience Research. 2005;79:216–23.

62. Basu A, Krishnamurthy S. Cellular Responses to Cisplatin-Induced DNA Damage. J Nucleic Acids. 2010;2010:201367.

63. Wei F, Hao P, Zhang X, Hu H, Jiang D, Yin A, et al. Etoposide-induced DNA damage affects multiple cellular pathways in addition to DNA damage response. Oncotarget. 2018;9:24122–39.

64. Grow EJ, Weaver BD, Smith CM, Guo J, Stein P, Shadle SC, et al. p53 convergently activates Dux/DUX4 in embryonic stem cells and in facioscapulohumeral muscular dystrophy cell models. Nature Genetics. 2021;53:1207–20.

65. Masteika IF, Sathya A, Homma S, Miller BM, Boyce FM, Miller JB. Downstream events initiated by expression of FSHD-associated DUX4: Studies of nucleocytoplasmic transport, γH2AX accumulation, and Bax/Bak-dependence. Biology Open. 2022;11.

66. Timinszky G, Till S, Hassa PO, Hothorn M, Kustatscher G, Nijmeijer B, et al. A macrodomain-containing histone rearranges chromatin upon sensing PARP1 activation. Nature Structural and Molecular Biology. 2009;16:923–9.

67. Kozlowski M, Corujo D, Hothorn M, Guberovic I, Mandemaker IK, Blessing C, et al. MacroH2A histone variants limit chromatin plasticity through two distinct mechanisms. EMBO reports [Internet]. 2018 [cited 2019 Sep 10];19.

68. Bensaude O. Inhibiting eukaryotic transcription. Transcription. 2011;2:103–8.

69. Frank L, Rippe K. Repetitive RNAs as Regulators of Chromatin-Associated Subcompartment Formation by Phase Separation. Journal of Molecular Biology. 2020;432:4270–86.

70. Chu HP, Cifuentes-Rojas C, Kesner B, Aeby E, Lee H goo, Wei C, et al. TERRA RNA Antagonizes ATRX and Protects Telomeres. Cell. 2017;170:86–101.e16.

71. Mondal T, Subhash S, Kanduri C. Chromatin RNA Immunoprecipitation (ChRIP). Methods Mol Biol. 2018;1689:65–76.

72. West JA, Davis CP, Sunwoo H, Simon MD, Sadreyev RI, Wang PI, et al. The long noncoding RNAs NEAT1 and MALAT1 bind active chromatin sites. Mol Cell. 2014;55:791–802.

73. Gaillard MC, Broucqsault N, Morere J, Laberthonnière C, Dion C, Badja C, et al. Analysis of the 4q35 chromatin organization reveals distinct long-range interactions in patients affected with Facio-Scapulo-Humeral Dystrophy. Scientific Reports. 2019;9.

74. Bodega B, Ramirez GDC, Grasser F, Cheli S, Brunelli S, Mora M, et al. Remodeling of the chromatin structure of the facioscapulohumeral muscular dystrophy (FSHD) locus and upregulation of FSHD-related gene 1 (FRG1) expression during human myogenic differentiation. BMC biology. 2009;7:41.

75. Ricci G, Zatz M, Tupler R. Facioscapulohumeral Muscular Dystrophy: More Complex than it Appears. Current Molecular Medicine. 2014;14:1052–68.

76. Mul K, Kinoshita J, Dawkins H, van Engelen B, Tupler R, Ferreira VA, et al. 225th ENMC international workshop:: A global FSHD registry framework, 18–20 November 2016, Heemskerk, The Netherlands. Neuromuscular Disorders. 2017;27:782–90.

77. Pastorello E, Cao M, Trevisan CP. Atypical onset in a series of 122 cases with FacioScapuloHumeral Muscular Dystrophy. Clin Neurol Neurosurg. 2012;114:230–4.

78. Nguyen K, Broucqsault N, Chaix C, Roche S, Robin JD, Vovan C, et al. Deciphering the complexity of the 4q and 10q subtelomeres by molecular combing in healthy individuals and patients with facioscapulohumeral dystrophy. Journal of Medical Genetics. 2019;56:590–601.

79. Ricci G, Ruggiero L, Vercelli L, Sera F, Nikolic A, Govi M, et al. A novel clinical tool to classify facioscapulohumeral muscular dystrophy phenotypes. J Neurol. 2016;263:1204–14.

80. Ricci G, Scionti I, Sera F, Govi M, D’Amico R, Frambolli I, et al. Large scale genotype-phenotype analyses indicate that novel prognostic tools are required for families with facioscapulohumeral muscular dystrophy. Brain. 2013;136:3408–17.

81. Ruggiero L, Mele F, Manganelli F, Bruzzese D, Ricci G, Vercelli L, et al. Phenotypic Variability Among Patients With D4Z4 Reduced Allele Facioscapulohumeral Muscular Dystrophy. JAMA network open. 2020;3:e204040.

82. Ricci G, Mele F, Govi M, Ruggiero L, Sera F, Vercelli L, et al. Large genotype–phenotype study in carriers of D4Z4 borderline alleles provides guidance for facioscapulohumeral muscular dystrophy diagnosis. Scientific Reports. 2020;10:1–12.

83. Vercelli L, Mele F, Ruggiero L, Sera F, Tripodi S, Ricci G, et al. A 5-year clinical follow-up study from the Italian National Registry for FSHD. Journal of Neurology. 2021;268:356–66.

84. Goto K, Nishino I, Hayashi YK. Very low penetrance in 85 Japanese families with facioscapulohumeral muscular dystrophy 1A. Journal of medical genetics. 2004;41:12e–12.

85. Sakellariou P, Kekou K, Fryssira H, Sofocleous C, Manta P, Panousopoulou A, et al. Mutation spectrum and phenotypic manifestation in FSHD Greek patients. Neuromuscular Disorders. 2012;22:339–49.

86. Zatz M, Marie SK, Passos-Bueno MR, Vainzof M, Campiotto S, Cerqueira A, et al. High proportion of new mutations and possible anticipation in Brazilian facioscapulohumeral muscular dystrophy families. American Journal of Human Genetics. 1995;56:99–105.

87. Salort-Campana E, Nguyen K, Bernard R, Jouve E, Solé G, Nadaj-Pakleza A, et al. Low penetrance in facioscapulohumeral muscular dystrophy type 1 with large pathological D4Z4 alleles: A cross-sectional multicenter study. Orphanet Journal of Rare Diseases. 2015;10:2.

88. Nakagawa M, Matsuzaki T, Higuchi I, Fukunaga H, Inui T, Nagamitsu S, et al. Facioscapulohumeral Muscular Dystrophy: Clinical Diversity and Genetic Abnormalities in Japanese Patients. Internal Medicine. 1997;36:333–9.

89. Wang Z, Qiu L, Lin M, Chen L, Zheng F, Lin L, et al. Prevalence and disease progression of genetically-confirmed facioscapulohumeral muscular dystrophy type 1 (FSHD1) in China between 2001 and 2020: a nationwide population-based study. Lancet Reg Health West Pac. 2022;18:100323.

90. He JJ, Lin XD, Lin F, Xu GR, Xu LQ, Hu W, et al. Clinical and genetic features of patients with facial-sparing facioscapulohumeral muscular dystrophy. European Journal of Neurology. 2018;25:356–64.

91. Lin F, Wang ZQ, Lin MT, Murong SX, Wang N. New insights into genotype-phenotype correlations in Chinese facioscapulohumeral muscular dystrophy: A retrospective analysis of 178 patients. Chinese Medical Journal. 2015;128:1707–13.

92. Ruggiero L, Mele F, Manganelli F, Bruzzese D, Ricci G, Vercelli L, et al. Phenotypic Variability Among Patients With D4Z4 Reduced Allele Facioscapulohumeral Muscular Dystrophy. JAMA Netw Open. 2020;3:e204040.

93. Rodolico C, Politano L, Portaro S, Murru S, Boccone L, Sera F, et al. Deletion of the Williams Beuren syndrome critical region unmasks facioscapulohumeral muscular dystrophy. European Journal of Paediatric Neurology. 2020;27:25–9.

94. Iodice R, Ugga L, Aruta F, Iovino A. Facioscapulohumeral muscular dystrophy (FSHD) and multiple sclerosis: a case report.

95. Lima da Silva G, Guimarães T, Pinto FJ, Brito D. A Unique Case of Type-1 Facioscapulohumeral Muscular Dystrophy and Sarcomeric Hypertrophic Cardiomyopathy. Rev Esp Cardiol. 2018;71:765–6.

96. Nauman F, Hussain MFA, Burakgazi AZ. The development of myasthenia gravis in a patient with facioscapulohumeral muscular dystrophy: case report and literature review. Neurology International [Internet]. 2019 [cited 2024 Mar 12];11.

97. Filippelli E, Barone S, Granata A, Nisticò R, Valentino P. A case of facioscapulohumeral muscular dystrophy and myasthenia gravis with positivity of anti-Ach receptor antibody: a fortuitous association? Neurol Sci. 2019;40:195–7.

98. Mammen AL, Casciola-Rosen L, Christopher-Stine L, Lloyd TE, Wagner KR. Myositis-specific autoantibodies are specific for myositis compared to genetic muscle disease. Neurol Neuroimmunol Neuroinflamm. 2015;2:e172.

99. Kocsis D, Herszényi L, Tóth M, Tulassay Z, Juhász M. Celiac Disease Associated with Facioscapulohumeral Muscular Dystrophy. International Journal of Celiac Disease. 2015;3:162–4.

100. Hangül C, Yücel OK, Toylu A, Uysal H, Berker Karaüzüm S. A Novel Coincidence: Essential Thrombocythemia with Facioscapulohumeral Muscular Dystrophy. Turk J Haematol. 2020;37:306–7.

101. Ziccone V, Rodolico C, Rizzo V, Tupler R, Buccafusca M, Toscano A. Facioscapulohumeral Muscular Dystrophy and Poliomyelitis followed by Multiple Sclerosis: A “triple trouble” case report and review of the literature on the association of MS and muscle disorders. Neuromuscul Disord. 2021;31:1179–85.

102. Hannah-Shmouni F, Al-Shahoumi R, Brady LI, Wu L, Frei J, Tarnopolsky MA. Dual molecular diagnoses in a neurometabolic specialty clinic. Am J Med Genet A. 2021;185:766–73.

103. Ottaviani A, Rival-Gervier S, Boussouar A, Foerster AM, Rondier D, Sacconi S, et al. The D4Z4 macrosatellite repeat acts as a CTCF and A-type lamins-dependent insulator in Facio-Scapulo-Humeral dystrophy. PLoS Genetics. 2009;5.

104. Arnoult N, Schluth-Bolard C, Letessier A, Drascovic I, Bouarich-Bourimi R, Campisi J, et al. Replication Timing of Human Telomeres Is Chromosome Arm–Specific, Influenced by Subtelomeric Structures and Connected to Nuclear Localization. PLoS Genet. 2010;6:e1000920.

105. Ottaviani A, Schluth-Bolard C, Gilson E, Magdinier F. D4Z4 as a prototype of CTCF and lamins-dependent insulator in human cells. Nucleus. 2010;1:30–6.

106. Campbell AE, Dyle MC, Albanese R, Matheny T, Sudheendran K, Cortázar MA, et al. Compromised nonsense-mediated RNA decay results in truncated RNA-binding protein production upon DUX4 expression. Cell Rep. 2023;42:112642.

107. Feng Q, Snider L, Jagannathan S, Tawil R, Van Der Maarel SM, Tapscott SJ, et al. A feedback loop between nonsense-mediated decay and the retrogene DUX4 in facioscapulohumeral muscular dystrophy Main text. 2015;4:4996.

108. Nickless A, Cheruiyot A, Flanagan KC, Piwnica-Worms D, Stewart SA, You Z. p38 MAPK inhibits nonsense-mediated RNA decay in response to persistent DNA damage in noncycling cells. J Biol Chem. 2017;292:15266–76.

109. Sato H, Singer RH. Cellular variability of nonsense-mediated mRNA decay. Nat Commun. 2021;12:7203.

110. de Koning APJ, Gu W, Castoe TA, Batzer MA, Pollock DD. Repetitive elements may comprise over Two-Thirds of the human genome. Copenhaver GP, editor. PLoS Genetics. 2011;7:e1002384.

111. Smit AF. Interspersed repeats and other mementos of transposable elements in mammalian genomes. Current Opinion in Genetics and Development. 1999;9:657–63.

112. Dumbovic G, Forcales SV, Perucho M. Emerging roles of macrosatellite repeats in genome organization and disease development. Epigenetics. 2017;12:515–26.

113. Brahmachary M, Guilmatre A, Quilez J, Hasson D, Borel C, Warburton P, et al. Digital Genotyping of Macrosatellites and Multicopy Genes Reveals Novel Biological Functions Associated with Copy Number Variation of Large Tandem Repeats. Orr HT, editor. PLoS Genetics. 2014;10:e1004418.

114. Stankiewicz P, Lupski JR. Structural variation in the human genome and its role in disease. Annual Review of Medicine. 2010;61:437–55.

115. Salsi V, Ferrari S, Ferraresi R, Cossarizza A, Grande A, Zappavigna V. HOXD13 Binds DNA Replication Origins To Promote Origin Licensing and Is Inhibited by Geminin. Molecular and Cellular Biology. 2009;29:5775–88.

