## Supplementary material for "Post-transcriptional RNA stabilization of telomere-proximal RNAs FRG2, DBET, D4Z4 at human 4q35 in response to genotoxic stress and D4Z4 macrosatellite repeat length": Salsi et al Supplemental Figures and legends

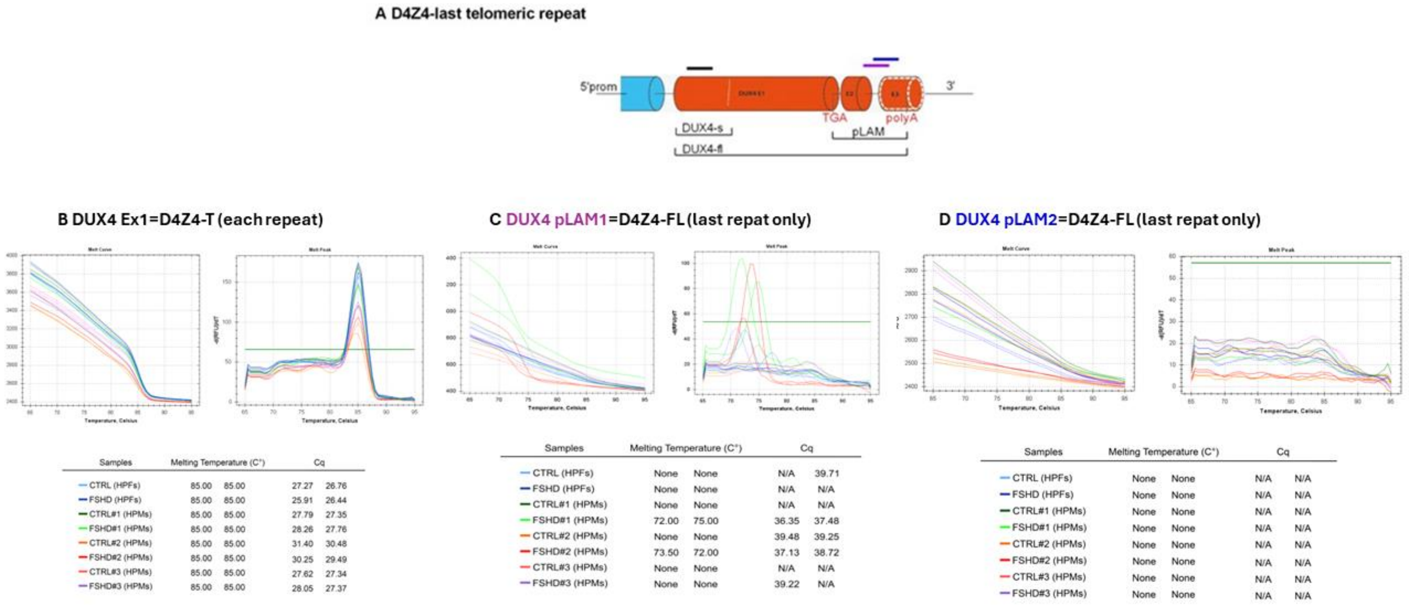


SF1 **Expression analysis of DUX4-T and DUX4FL.** A) Schematic representation of the last D4Z4 unit, the adjacent pLAM region and the distal exons. The DUX4 ORF is contained in the first exon in each repeat. The pLAM sequence is only present in the DUX4-FL transcript. Primers used in for qPCR are indicated in different colors. B-D) qPCR reports of DUX-T and DUX-FL amplification by using the indicated primer pairs, melting cure graphs are visible in the upper part of each graph, melting curve and quantification cycle ($C_q$) reports are indicated in the table below for human primary myoblasts (HPMs) and human primary trophoblast cells (here HPFs). For DUX-FL we used two different primer sets (pLAM1, purple and pLAM2 blue) used in previous works [48,49].

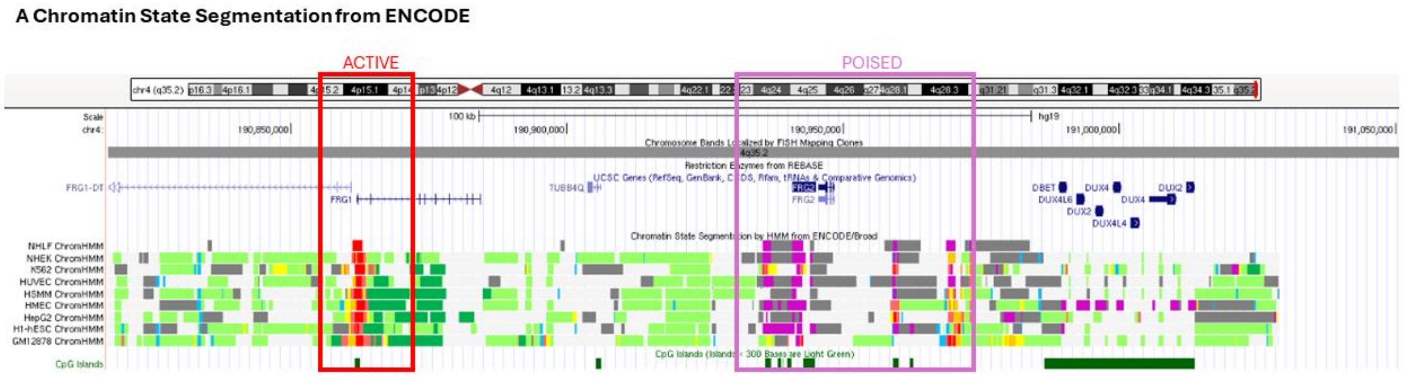


SF2 **Chromatin State Segmentation by HMM from ENCODE/Broad; Hg19chr4:190,793,373-191,038,972.**

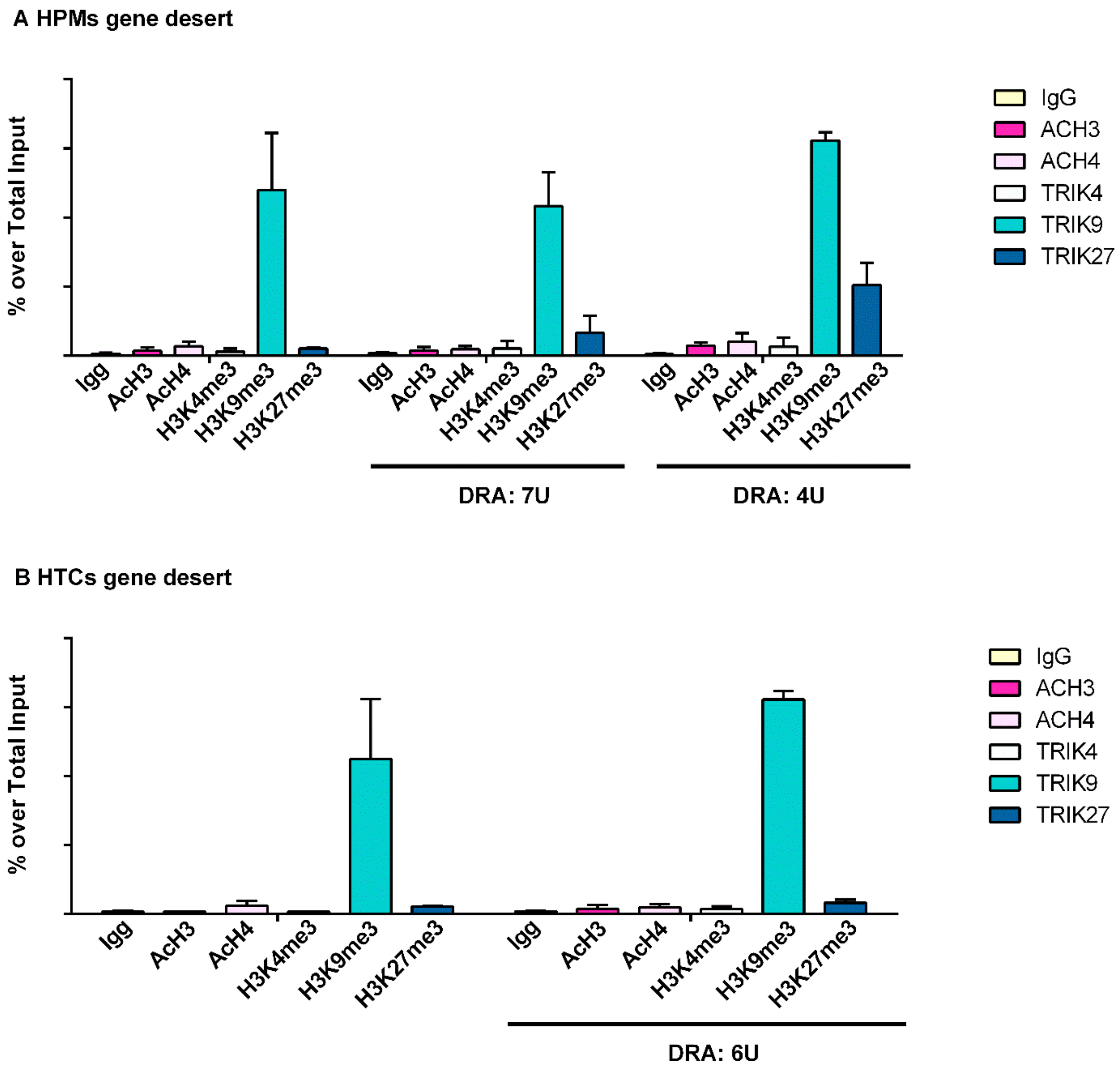


SF3 **Epigenetic status of 4q35-associated gene desert region**. Chromatin immunoprecipitation assays (ChIP) conducted in (A) HPMs and HTCs (B) carrying a normal sized (control) and reduced (DRA) D4Z4 alleles. Antibodies directed to H3K4me3, H3K9me3, H3K27me3 and pan-acetylated Histone 3 and 4 (AcH3 and AcH4) were used, followed by qPCR amplification using primers described in Fig1.A.

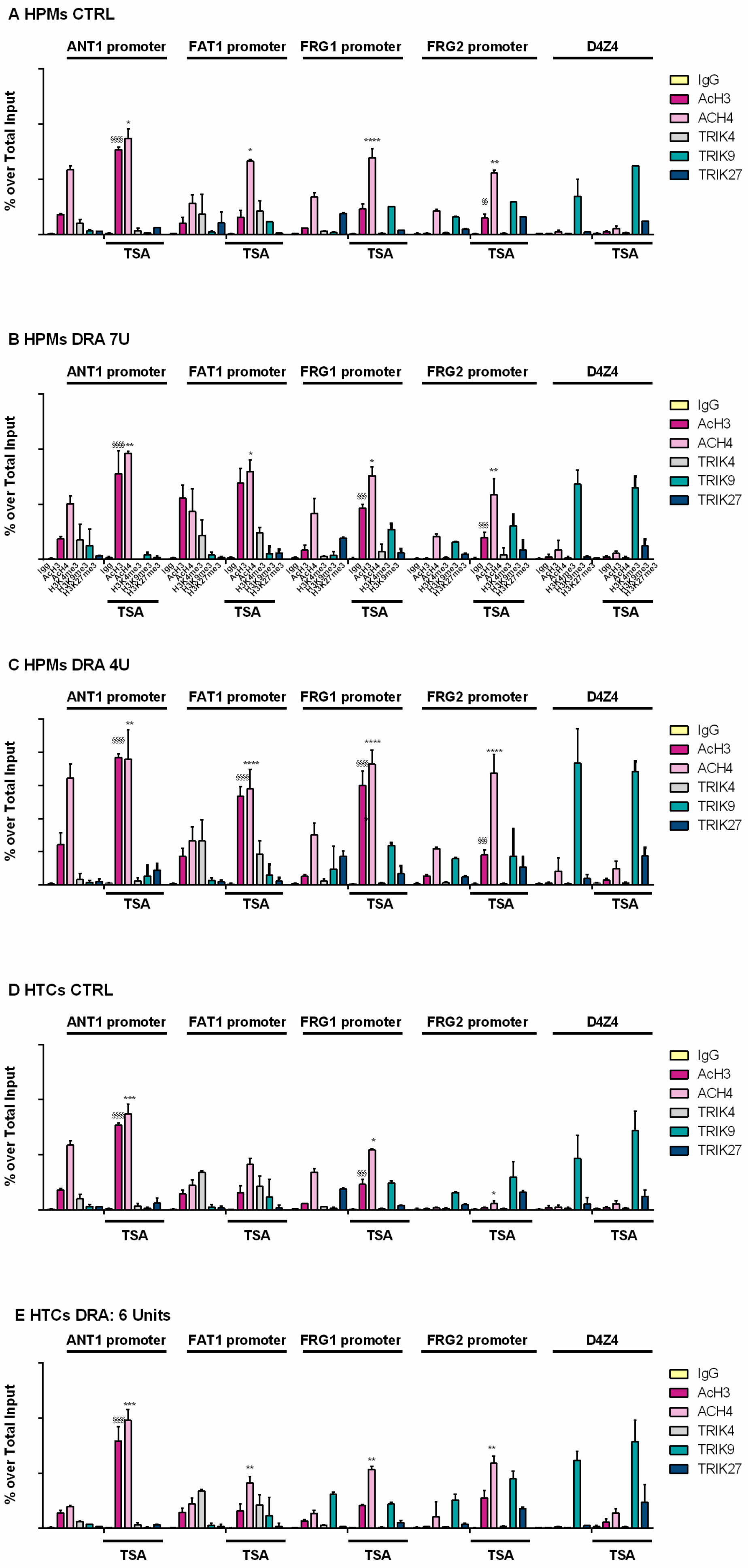
A HPMs CTRL
B HPMs DRA 7U
C HPMs DRA 4U
D HTCs CTRL
E HTCs DRA: 6 Units
ANT1 promoter
FAT1 promoter
FRG1 promoter
FRG2 promoter
D4Z4
% over Total Input
TSA
IgG
AcH3
ACH4
TRIK4
TRIK9
TRIK27

SF4. G-H) **Chromatin immunoprecipitation assays (ChIP)** conducted in HPMs (A-C) and HTCs (D-E) carrying a normal sized (control) and reduced (DRA) D4Z4 alleles (B,C,E) and treated or not with TSA. Antibodies directed to H3K4me3, H3K9me3, H3K27me3 and pan-acetylated Histone 3 and 4 (AcH3 and AcH4) were used, followed by qPCR amplification using primers described in Fig1.A. Anova statistical test with multiple comparison was performed (*0.05<p value<0.01; ** 0.01<p value<0.001; *** 0.001<p value<0.0001; **** P value<0.0001): $ (dollar symbol), and * (asterisk). Dollar symbols and asterisks indicate the statistical significance of data obtained in TSA treated cells in respect to the same antibody enrichment in not treated cells. 4q35 regions amplified in qPCR following ChIP are indicated in the upper part of each graph.

**Human Primary Trophooblasts (HTCs)**

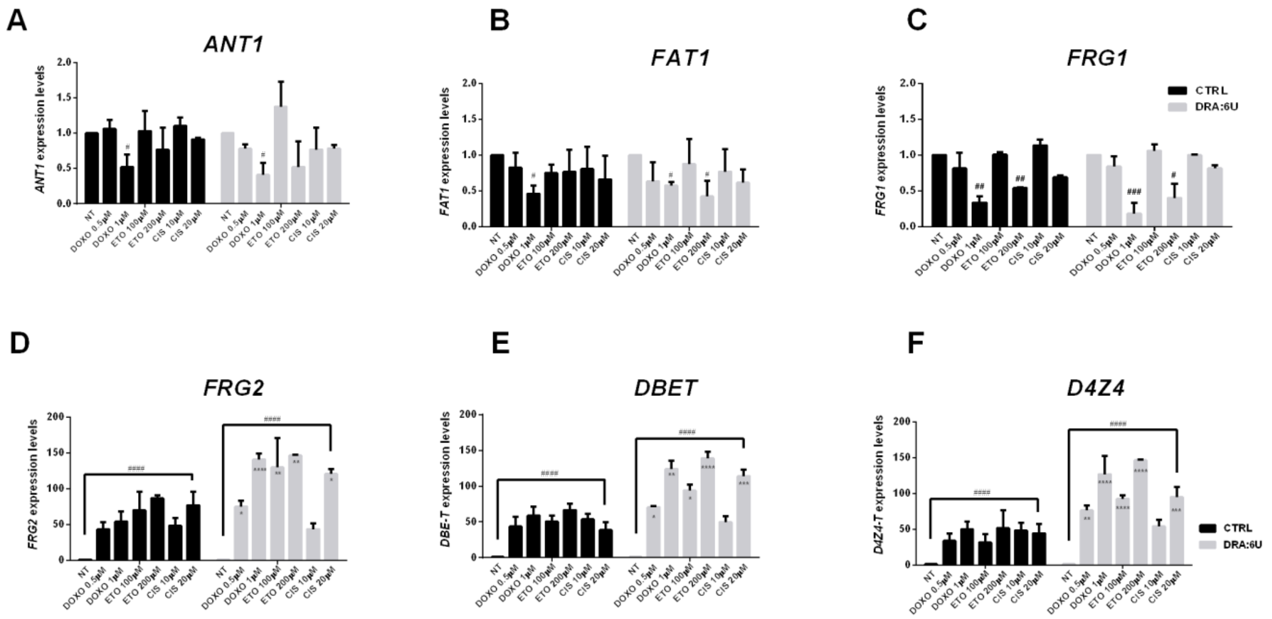


SF5 **Control HTCs and HTCs bearing 6U D4Z4 array were untreated or treated with genotoxic drugs**: Doxorubicin (DOXO), Etoposide (ETO) and Cisplatin (CIS), at the reported concentrations. Expression data of *ANT1* (A), *FAT1* (B), *FRG1* (C), *FRG2* (D), *DBE-T* (E) and *D4Z4-T* (F) was evaluated 24h after treatments and normalized over *RPLP0* reference gene levels. Error bars represent standard deviation values for three independent replicates.

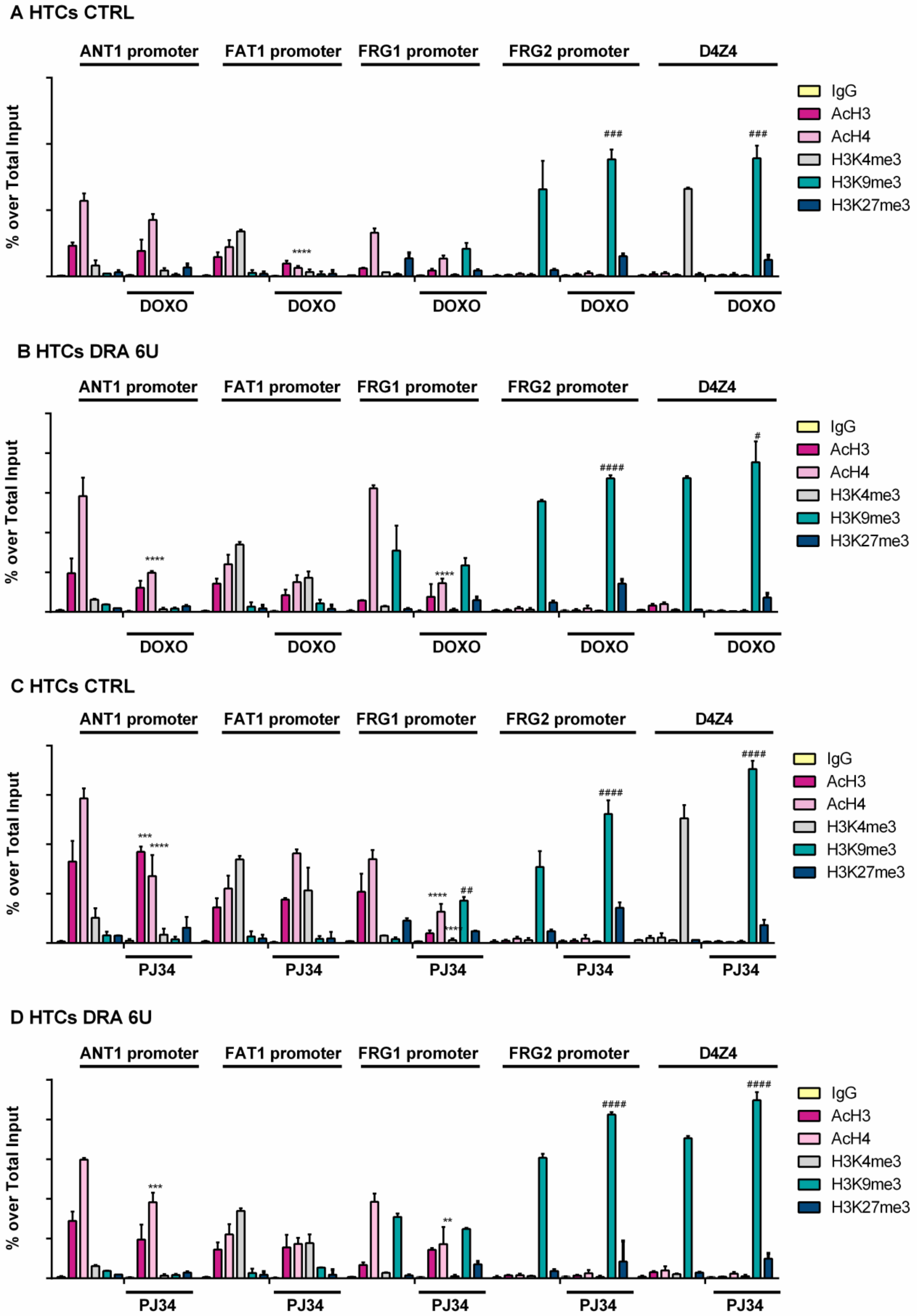


**Supplemental_ Fig_S6**

SF6 A-D )**Chromatin immunoprecipitation assays (ChIP)** conducted in control and DRA HTCs treated or not with Doxorubicin (A,B) or PJ34 (C,D). Antibodies directed to H3K4me3, H3K9me3, H3K27me3 and pan-acetylated Histone 3 and 4 (AcH3 and AcH4) were used, followed by qPCR amplification using primers described in Fig1.A. Anova statistical test with multiple comparison was performed (*0.05<p value<0.01; ** 0.01<p value<0.001; *** 0.001<p value<0.0001; **** P value<0.0001). Different symbols: * (asterisk) and # (hashtag) refer to different antibodies used in ChIP experiments (*=AcH4; #=H3K9me3 to show the statistical significance of data obtained in treated cells in respect to the same in not treated cells). . Error bars represent standard deviation values for three independent replicates.
